# GTSE1 tunes microtubule stability for chromosome alignment and segregation through MCAK inhibition

**DOI:** 10.1101/067827

**Authors:** Shweta Bendre, Arnaud Rondelet, Conrad Hall, Nadine Schmidt, Yu-Chih Lin, Gary J. Brouhard, Alexander W. Bird

## Abstract

The dynamic regulation of microtubules during mitosis is critical for accurate chromosome segregation and genome stability. Cancer cell lines with hyperstabilized kinetochore microtubules have increased segregation errors and elevated chromosomal instability (CIN), but the genetic defects responsible remain largely unknown. The microtubule depolymerase MCAK can influence CIN through its impact on microtubule stability, but how its potent activity is controlled in cells remains unclear. Here we show that GTSE1, a protein found overexpressed in aneuploid cancer cell lines and tumours, regulates microtubule stability during mitosis by inhibiting MCAK microtubule depolymerase activity. Cells lacking GTSE1 have defects in chromosome alignment and spindle positioning due to microtubule instability caused by excess MCAK activity. Reducing GTSE1 levels in CIN cancer cell lines reduces chromosome missegregation defects, while artificially inducing GTSE1 levels in chromosomally stable cells elevates chromosome missegregation and CIN. Thus GTSE1 inhibition of MCAK activity regulates the balance of microtubule stability that determines the fidelity of chromosome alignment, segregation, and chromosomal stability.

## Introduction

The precise regulation of microtubule (MT) dynamics is essential to the accurate execution of mitosis and faithful segregation of chromosomes. Defects in the regulation of MT stability and dynamics can result in errors in spindle positioning and chromosome segregation, two processes found defective in cancers (Gordon et al., 2012; Noatynska et al., 2012). Persistent errors in chromosome segregation lead to chromosomal instability (CIN), the increased rate of gain or loss of chromosomes within a cell population. CIN is present in most solid tumours, and recent evidence suggests CIN plays a causal role in tumorigenesis (Schvartzman et al., 2010). The genetic and molecular defects that lead to CIN in tumours, however, remain largely unknown.

In several cancer cell lines with CIN, kinetochore-MT attachments are hyperstabilized (Bakhoum et al., 2009a). This leads to an increased frequency of chromosome missegregation, and ultimately to CIN, due to a reduced ability of cells to correct erroneous kinetochore-MT attachments, in particular merotelic attachments, where one kinetochore is connected to microtubules from both spindle poles (Bakhoum et al., 2009a; b). Cells must therefore be able to precisely regulate MT dynamics so that kinetochore MTs are dynamic enough to correct erroneous attachments, yet stable enough to efficiently capture and align chromosomes (Bakhoum et al., 2009a; b). The regulatory mechanisms by which cells are able to maintain this balance and avoid CIN remain unclear.

A major direct regulator of MT stability is the kinesin-13 MT depolymerase MCAK/Kif2C. *In vitro*, MCAK has extremely potent depolymerase activity (Desai et al., 1999; Hunter et al., 2003; Helenius et al., 2006). In cells, reduction of MCAK activity leads to an increase in microtubule polymer (Rankin and Wordeman, 2010; Rizk et al., 2009). Microtubule-kinetochore attachments are also hyperstabilized, leading to defects in correcting merotelic attachments and in chromosome segregation (Bakhoum et al., 2009a; Maney et al., 1998; Kline-Smith et al., 2004). Excessive MCAK activity induced by overexpression of MCAK leads to loss of MT stability throughout the cell and defects in the capture and alignment of chromosomes (Maney et al., 1998; Moore and Wordeman, 2004; Zhang et al., 2011). MCAK MT depolymerase activity must therefore be precisely controlled in time and cellular space to ensure both chromosome alignment and segregation and to avoid CIN. While interest in MCAK regulation has led to the identification of proteins that enhance or counteract MCAK activity in cells (Cross and Powers, 2011; Ohi et al., 2003; Jiang et al., 2009; Meunier and Vernos, 2011), only NuSAP has been recently reported to attenuate MCAK activity via direct interaction (Li et al., 2015). *In vitro* studies of MCAK have uncovered potential mechanisms by which intramolecular rearrangements of MCAK can determine MT depolymerase activity (Ems-McClung et al., 2013; Talapatra et al., 2015; Burns et al., 2014). Based on this knowledge, proposed mechanisms for direct regulation of MCAK activity in cells have thus largely relied on intramolecular rearrangements induced from interaction with microtubules, nucleotide exchange, and phosphorylation by mitotic kinases (Ems-McClung et al., 2013; Talapatra et al., 2015; Burns et al., 2014; Cooper et al., 2009).

Because MCAK activity affects kinetochore MT stability, its deregulation may impact CIN. Indeed, artificially destabilizing kinetochore MTs in CIN lines by overexpressing MCAK reduces chromosome missegregation and CIN (Bakhoum et al., 2009b). While these key experiments point to hyperstability of kinetochore MTs in cancer cell lines as a direct cause of CIN, they do not resolve the molecular genetic origin of this defect, as MCAK protein levels are not generally downregulated in cancer cell lines or tumours (Bakhoum et al., 2009a; Sanhaji et al., 2011). Therefore, investigation into cellular regulation of MCAK activity, as well as the molecular basis of kinetochore MT hyperstabilization in cancer cells, is highly desirable.

GTSE1 is a microtubule-associated and EB1-dependent plus-end tracking protein (Monte et al., 2000; Scolz et al., 2012). In interphase, recruitment of GTSE1 to growing microtubule plus-ends by EB1 is required for cell migration. During mitosis, GTSE1 is heavily phosphorylated and no longer tracks microtubule plus-ends nor interacts directly with the microtubule lattice, but localizes to the mitotic spindle in a TACC3-dependent manner (Scolz et al., 2012; Hubner et al., 2010). GTSE1 expression levels are upregulated in several tumour types and cancer cell lines, and correlate with tumour grade and poor clinical outcome in breast cancers (Scolz et al., 2012), although it remains unknown whether GTSE1 overexpression facilitates tumorigenesis.

Here we show that GTSE1 stabilizes MTs during mitosis to facilitate chromosome alignment by suppressing MCAK MT depolymerase activity. GTSE1 interacts with MCAK and inhibits its activity *in vitro*, thus extending our understanding of the mechanisms controlling MCAK activity. Perturbing the balance of the GTSE1-MCAK relationship in cancer cell lines influences CIN though its effect on MT stabilization: increasing GTSE1 expression in diploid cell lines leads to chromosome segregation defects and induces CIN, while depletion or knockout of the high GTSE1 levels in CIN cell lines reduces chromosome segregation defects in an MCAK-dependent manner. Thus, GTSE1 tunes MT stability in mitosis to ensure both chromosome alignment and accurate segregation by suppressing MCAK MT depolymerase activity.

## Results

### GTSE1 stabilizes microtubules in mitosis and promotes correct spindle orientation

To investigate the role of GTSE1 during mitosis, we first reduced GTSE1 protein levels by RNAi in human U2OS cells and analyzed them by immunofluorescence (Fig. 1 A and S1 A). GTSE1-depleted mitotic cells displayed a loss of microtubule stability as observed by immunofluorescence labeling of tubulin. In more than 50% of cells depleted of GTSE1, astral microtubules appeared absent or dramatically shortened in both prometaphase and metaphase cells (Fig. 1, A and B), and microtubule density within the inner spindle was significantly reduced (Fig. 1 C). We quantified the defect in astral MT stability via three additional analyses. First, we measured the length in three dimensions of the 10 longest astral MTs visible per cell from tubulin immunfluorescence. The length of the few visible astral MTs remaining after GTSE1 RNAi was less than half that of control cells (Fig 1 D). Second, we calculated the average length of all astral MTs per cell by calculating the average distance between growing astral MT plus-ends and the centrosome via EB1 protein immunofluorescence, and again found a significant reduction (Fig 1E). Finally, we calculated the total number of growing astral MTs from EB1 staining, and found a nearly 50% reduction following GTSE1 RNAi (Fig 1F).

**Figure 1.**
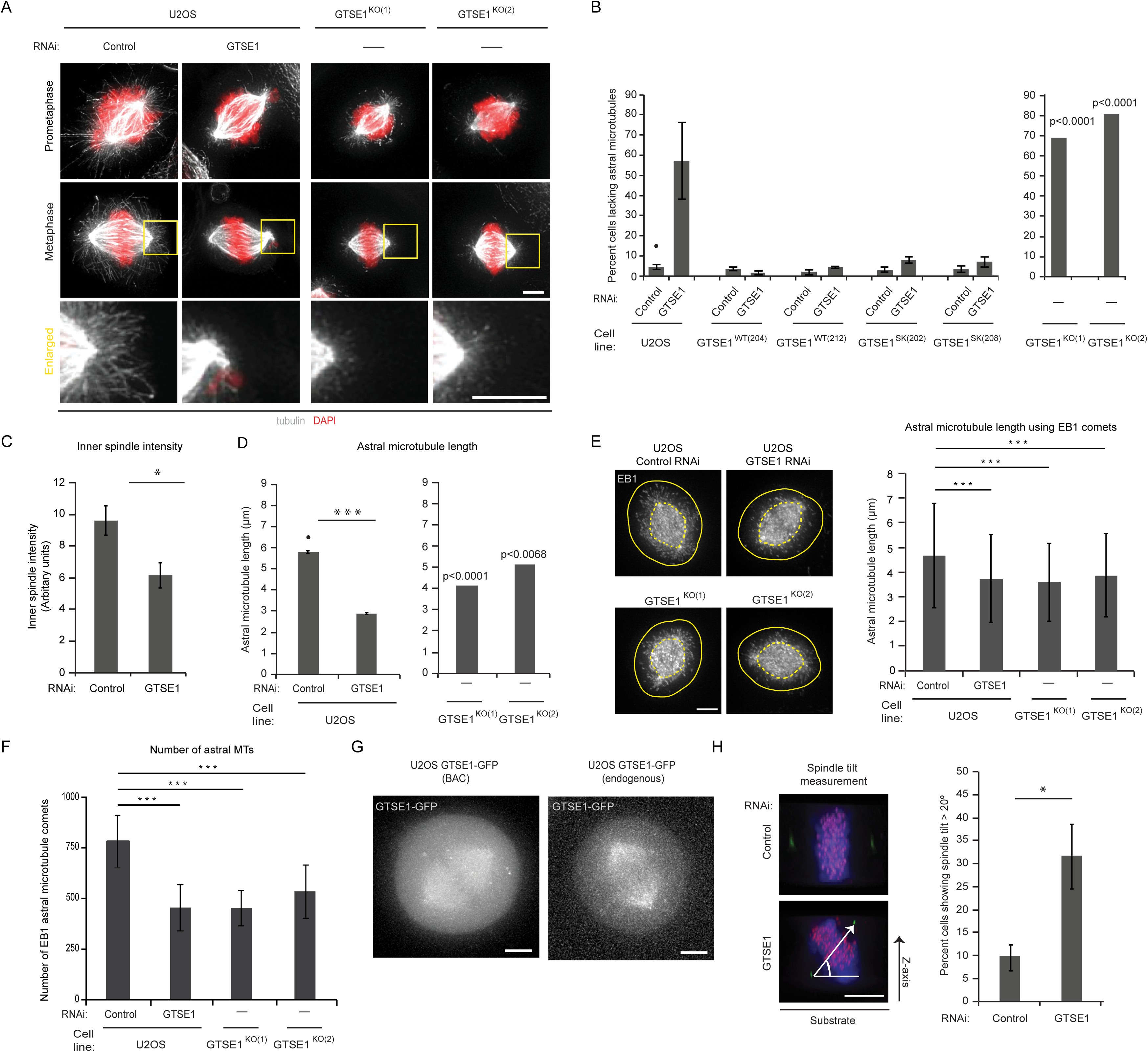
GTSE1 stabilizes microtubules in mitosis and promotes correct spindle orientation. (A) Immunofluorescence images of U2OS cells after control RNAi, GTSE1 RNAi, and stable knockout of GTSE1 stained for DNA (DAPI) and microtubules (tubulin) showing less astral microtubules following reduced GTSE1 levels. (B) Quantification of the percent cells lacking astral microtubules from immunofluorescence analysis as shown in A. The left bar chart shows control and GTSE1 RNAi in U2OS cells, two stable U2OS cell clones expressing RNAi-resistantGTSE1 (GTSE1 ^WT(204)^ and GTSE1 ^WT(212)^), and two stable U2OS cell clones expressing RNAi-resistant GTSE1 mutated to abolish interaction with EB1 (GTSE1 ^Sk(202)^ andGTSE1 ^Sk(208)^). n > 150 cells over 3 experiments per condition. The right bar chart shows two independent U2OS GTSE1 knockout clones. P-values were obtained from a chi-squared test comparing to the U2OS control condition (marked with •). n > 100cells (C) Quantification of “inner-spindle” tubulin fluorescence intensity from fixedU2OS cells stained for alpha-tubulin following control or GTSE1 RNAi. n ≥ 19 per experiment per condition; 3 experiments. (D) Quantification of the average length of astral microtubules in three dimensions from immunofluorescence analysis as shown in A. 10 astral MTs were measured per cell. n = 10 cells per experiment per condition; 3 experiments (E) Immunofluorescence images of U2OS cells after control RNAi, GTSE1 RNAi, and stable knockout of GTSE1 stained for EB1 showing less EB1 astral microtubule comets following GTSE1 RNAi. Bar chart shows quantification of astral microtubule length using the position of EB1 comets with respect to the centrosome. n > 5800 astrals from a single experiment. Error bars represent standard deviation. P-values were obtained using a Kruskal-Wallis test followed by Conover-Iman test. (F) Quantification of the average number of astral microtubules per cell obtained by quantifying the number of EB1 comets in U2OS cells after control or GTSE1 RNAi and in GTSE1 ^KO(1)^ and GTSE1 ^KO(2)^ cells. n ≥ 13 cells per condition from a single experiment. Error bars represent standard deviation. P-values were obtained using a Anova and a Tukey’s test (G) Live-cell fluorescence images ofmetaphase U2OS cells expressing either bacterial artificial chromosome-expressed GTSE1-GFP (GTSE1 ^WT(212)^) or endogenously tagged GTSE1-GFP. Both constructs localize to the spindle. (H) Analysis of spindle orientation. Images of mitotic cells viewed from the side and stained for DNA (blue), kinetochores (red) and centrioles (green) depict a cell with normal spindle alignment parallel to the substrate (top image), and a cell with defective orientation (bottom image). The angle of spindle tilt was calculated by determining the angle between the substrate and a line connecting both centrosomes, as depicted. Quantification of the percent of metaphase cells with a spindle tilt angle greater than 20 degrees is shown. n > 140 cells over 3 experiments per condition. All scale bars represent 5 microns and all error bars when not specified represent standard error of the mean. p≤0.05 *, p≤0.01 **,p≤0.001 ***

To verify that the loss of MT stability following RNAi was specific to depletion of GTSE1, we depleted GTSE1 in two independent and clonal cell lines containing stably integrated RNAi-resistant, GFP-tagged GTSE1 genes harbored on bacterial artificial chromosomes (BACs)(GTSE1^WT(204)^ and GTSE1^WT(212)^)(Scolz et al., 2012; Bird et al., 2011). Following depletion of endogenous GTSE1, GTSE1-GFP localized to the mitotic spindle, which we also observed upon tagging of endogenous GTSE1 with GFP via Cas9/CRISPR-mediated homologous recombination (Cong et al., 2013; Ran et al., 2013) (Figs. 1 G and S2 A). RNAi-resistant GTSE1-GFP expressing cells maintained astral MTs after GTSE1 RNAi, confirming specificity (Fig. 1 B and S1 B).

To further confirm the role for GTSE1 in microtubule stability, we knocked out the genomic copies of the GTSE1 gene in U2OS cells via Cas9/CRISPR nuclease targeting (Cong et al., 2013; Ran et al., 2013). Interestingly, we were able to isolate viable clonal GTSE1 knockout cell lines, and two independently-constructed knockout clones were subsequently analyzed (GTSE1^KO(1)^ and GTSE1^KO(2)^ (Fig S2, B-D). Both GTSE1 knockout cell lines displayed a significant destabilization of astral microtubules in mitosis (Fig. 1, A, B, D, E and F). Although spindle microtubule stability was compromised, neither knockout nor RNAi depletion of GTSE1 led to a change in spindle length (Fig S2 E).

We previously found that the interaction between GTSE1 and EB1 is required for GTSE1’s function in cell migration and microtubule-dependent focal adhesion disassembly (Scolz et al., 2012). To determine whether this interaction was important for the role of GTSE1 in mitotic MT stability, we analyzed cells expressing RNAi-resistant, BAC-based GTSE1 that does not interact with EB1 due to mutation of conserved EB-binding (SxIP) motifs (Scolz et al., 2012). After depletion of endogenous GTSE1, SxIP-mutated GTSE1 was able to maintain astral stability similar to the wild-type GTSE1 gene (Figs. 1 B and S1 B). Thus, the interaction between GTSE1 and EB1 is not important for MT stability in prometaphase and metaphase.

GTSE1-depleted cells lacking astral microtubules often displayed a defect in the positioning of the spindle within the cell, consistent with known roles of astral MT-cortex interactions in establishing spindle orientation (Pearson and Bloom, 2004). We quantified this defect in spindle orientation by measuring the angle of the spindle relative to the coverslip surface, and found that cells depleted of GTSE1 more frequently failed to align the spindle axis within 20 degrees relative to the surface (Fig. 1 H). Thus, GTSE1 is necessary for the stabilization of microtubules in mitosis and proper spindle orientation.

### GTSE1 is required for efficient chromosome alignment

To determine whether GTSE1 plays a role in mitotic progression and chromosome segregation, we imaged live U2OS cells expressing histone mH2A.Z-mCherry progressing through mitosis following RNAi (Figs 2 A and S1 C, Videos 1 and 2). GTSE1-depleted cells took significantly longer than control-depleted cells to align all of their chromosomes and enter anaphase (control RNAi: 28.4 ± 20.5 min; GTSE1 RNAi: 42.5 ± 36.3 min)(Fig 2 B). To quantify the chromosome alignment defect, we fixed cells after RNAi and determined the percentage of cells containing misaligned chromosomes but otherwise displaying a metaphase-like morphology (Fig. 2, C and D). Approximately 40% of GTSE1-depleted cells contained misaligned chromosomes, and this defect was rescued by expression of a wildtype RNAi-resistant GTSE1-GFP transgene, as well as the SxIP-mutated transgene, confirming the specificity of GTSE1 RNAi (Fig 2D). Misaligned chromosomes were also significantly increased in both GTSE1^KO^ cell lines, although to a lesser extent, suggesting these cell lines may be adapted through selection of an (epi)genetic alteration that increases fidelity of chromosome segregation and viability in a GTSE1-knockout background. We analyzed protein levels of several mitotic regulators related to microtubule dynamics and GTSE1 function in these knockout cells, but did not see differences to wild-type cells (Fig S2 B,F-H).

**Figure 2.**
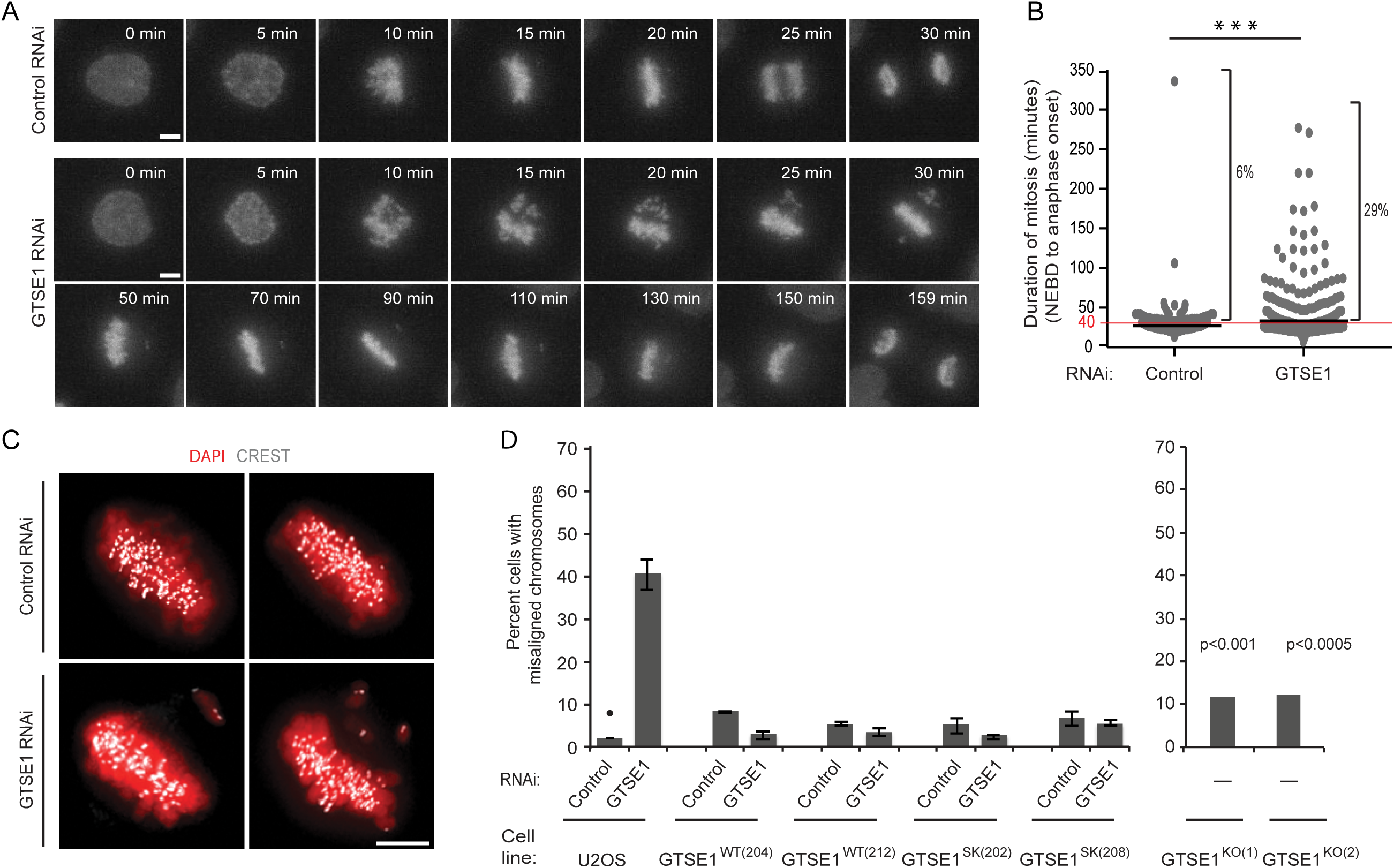
GTSE1 is required for efficient chromosome alignment. (A) Still frames ofmitoses from time-lapse movies (Supplemental Videos 1 and 2) of U2OS cells expressing histone H2AFZ-mCherry after control (top row) or GTSE1 RNAi (bottom two rows). GTSE1-depleted cells require a longer time to align all chromosomes and enter anaphase. (B) Mitotic duration (nuclear envelope breakdown (NEBD) toanaphase onset) of individual cells is plotted from the analysis of movies of control orGTSE1 RNAi-treated U2OS histone H2AFZ-mCherry cells as are shown in A. GTSE1-depleted cells have a longer average duration of mitosis (control RNAi: 28.4 ± 20.5 min, n=269; GTSE1 RNAi: 42.5 ± 36.3 min, n = 295; black bars represent the average)and a higher percentage of cells with mitotic duration longer than 40 minutes (5.9% of control depleted cells versus 29.2% of GTSE1-depleted cells). (C)Immunofluorescence images of chromosomes (red) and kinetochores (white) infixed U2OS cells after control or GTSE1 RNAi. (D) Quantification of the percent cellswith misaligned chromosomes from immunofluorescence analysis as shown in C. Theleft bar chart shows control or GTSE1 RNAi of U2OS cells, two stable U2OS cell clones expressing RNAi-resistant GTSE1 (GTSE1 ^WT(204)^ and GTSE1 ^WT(212)^), and two stableU2OS cell clones expressing RNAi-resistant GTSE1 mutated to abolish interaction with EB1 (GTSE1 ^Sk(0)^ and GTSE1 ^Sk(208)^). n > 150 cells over 3 experiments per condition. The right bar chart shows two independent U2OS GTSE1 knockout clones.P-values were obtained from a chi-squared test comparing to the U2OS controlcondition (marked with •). n > 100 cells. p≤0.05 *, p≤0.01 **, p≤0.001 ***

### GTSE1 stabilizes kinetochore-MT attachment

The defect in chromosome alignment and delay in anaphase onset in GTSE1-depleted cells indicated that the spindle assembly checkpoint was active in these cells, and suggested that they may be compromised for their ability to establish proper and stable kinetochore-MT attachments. To more specifically analyze this, we first quantified the abundance of MAD1, which is recruited to kinetochores that do not have proper kinetochore-MT attachments (Howell et al., 2004), in metaphase cells with aligned chromosomes. Indeed, GTSE1-depleted metaphase cells displayed increased kinetochore localization of MAD1 (Fig. 3 A). The delays in chromosome alignment and MAD1 persistence at kinetochores following GTSE1 depletion could arise from destabilization of MTs and kinetochore-MT attachments. To determine whether GTSE1 was required specifically for the stability of kinetochore MTs, we first tested whether there was a reduction in the cold-stable kinetochore MT population of metaphase cells following GTSE1 depletion. Indeed, GTSE1-depleted cells had a significant reduction of cold-stable kinetochore MTs as compared to control-treated cells (Fig. 3 B). To quantify kinetochore MT stability, we then assayed kinetochore MT turnover by measuring loss of fluorescence after photoactivation in metaphase U2OS cells expressing photoactivatable (PA) GFP-tubulin (Zhai et al., 1995; Bakhoum et al., 2009b). Analysis of fluorescence dissipation after photoactivation indicated that cells depleted of GTSE1 showed a decrease in the half-life of kinetochore MTs as compared to control RNAi (Figs. 3 C, S1 D, and S3 A), indicating a reduction in the stability of kinetochore-MT attachments. Consistent with a role in stabilizing kinetochore MTs, we observed that GTSE1 localized to kinetochore MTs, as visualized by removing non-kinetochore MTs by cold treatment and staining for GTSE1 and kinetochores (Fig 3D, Video 3 and 4). Together, these results implicate GTSE1 as an important regulator of microtubule stability in mitosis, necessary to stabilize kinetochore-MT attachment and promote efficient alignment of chromosomes.

**Figure 3.**
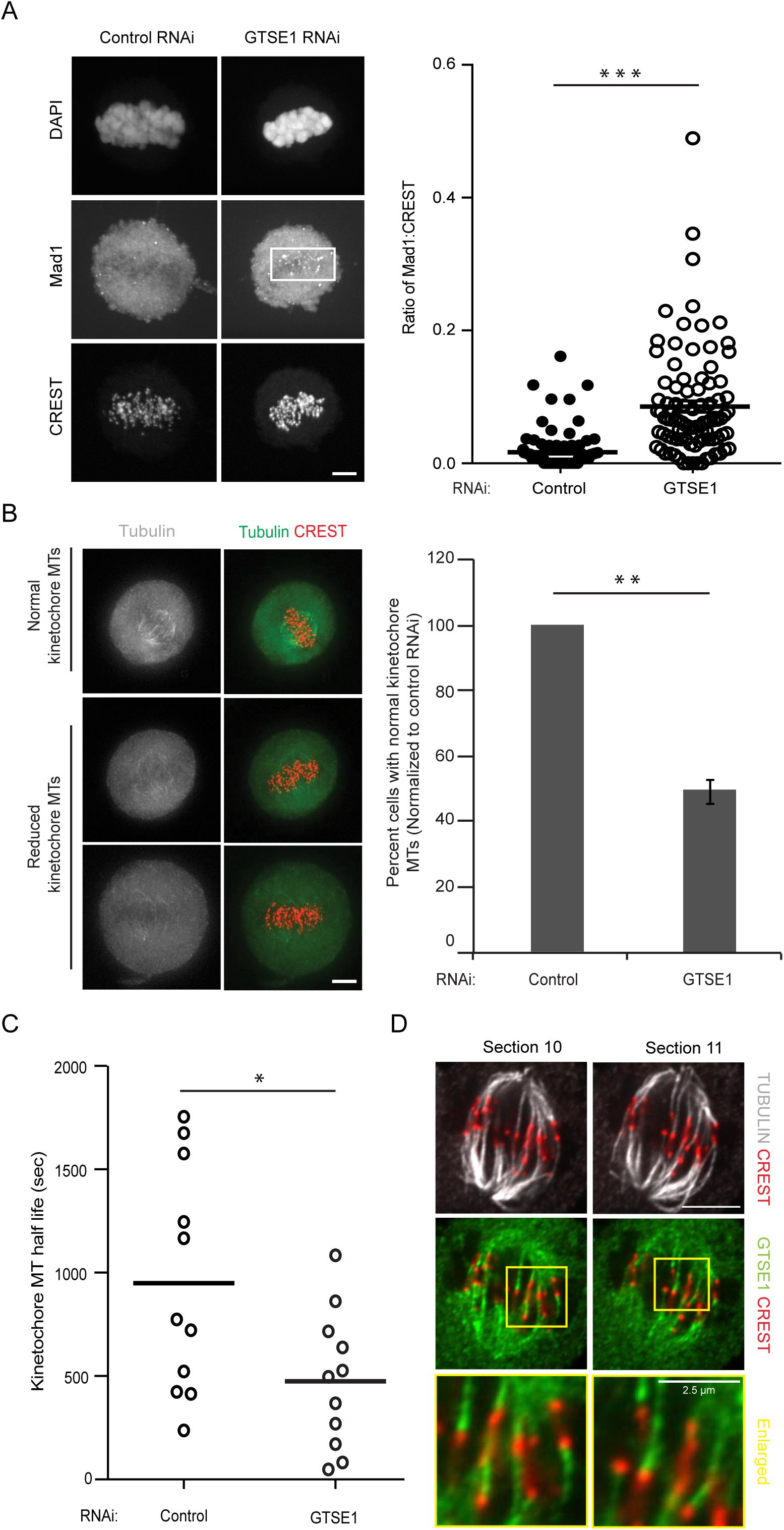
GTSE1 localizes to kinetochore microtubules and stabilizes kinetochore-microtubule attachments. (A) Representative immunofluorescence images of metaphase U2OS cells stained with DAPI (DNA), MAD1 and CREST showing MAD1 accumulation on kinetochores after GTSE1 RNAi. Cells were blocked using Cdk1inhibitor (RO-3306) and released for 1 hour in normal media to enrich the number of metaphase mitotic cells. n > 90 cells from three independent experiments. (B) Fewer GTSE1-depleted cells contain cold-stable microtubules than control-depleted cells. Images show mitotic cells fixed following cold treatment and stained for tubulin and kinetochores. The bar chart shows the proportion of cells containing a full complement of cold-stable microtubules following cold treatment, after control cells were normalized to 100%. The cells were blocked using Cdk1 inhibitor (RO-3306)and released for 1 hour in normal media to enrich the number of metaphase mitotic cells. n > 100 cells per experiment; 3 experiments per condition. (C) Graph showing kinetochore microtubule half-life in control or GTSE1-depleted U2OS cells expressing photoactivatable (PA) GFP-tubulin (see Fig. S 3). Each circle represents the kinetochore microtubule half-life of a single cell; bars represent the average. n = 11cells over 3 experiments per condition. (D) Representative immunofluorescence images (Slice 10 and 11 from Supplementary Movie 3 and 4) showing GTSE1 decorating K-fibers in U2OS cells expressing BAC based GTSE1-LAP following cold treatment. The cells were stained for GTSE1, kinetochores (CREST) and microtubules (tubulin). All error bars represent standard error of the mean. All scale bars represent 5 microns, unless otherwise stated. p≤0.05 *, p≤0.01 **, p≤0.001 ***

### Mitotic defects following GTSE1-depletion are dependent on the activity of MCAK

We next asked the mechanism by which GTSE1 promotes microtubule stability in mitosis. GTSE1 is highly phosphorylated when cells enter mitosis, and does not associate directly with growing microtubule tips nor the microtubule lattice (Scolz et al., 2012) (and unpublished results), indicating that the stabilizing effect of GTSE1 on microtubules in mitosis is mediated through other protein interactors. We previously showed that GTSE1 interacts with a complex containing TACC3 (Hubner et al., 2010), which has also been implicated in controlling microtubule stability (Kinoshita, 2005; Lin et al., 2010; Booth et al., 2011; Cheeseman et al., 2013; Nixon et al., 2015). We did not, however, detect a significant change in TACC3 localization to microtubules in GTSE1-depleted cells nor GTSE1 knockout cell lines (Fig S3 B), consistent with our earlier report (Hubner et al., 2010). Analysis of mitotic interactors of GTSE1 from our previous immunoprecipitation and mass spectrometry results (Hubner et al., 2010) revealed that the microtubule depolymerase MCAK (Kif2C) was also consistently enriched across repeated experiments. We confirmed this interaction in cells by immunoprecipitating either endogenous or stably BAC-expressed GFP-tagged GTSE1 from mitotic cells and probing with antibodies against MCAK (Fig. 4, A and B).

**Figure 4.**
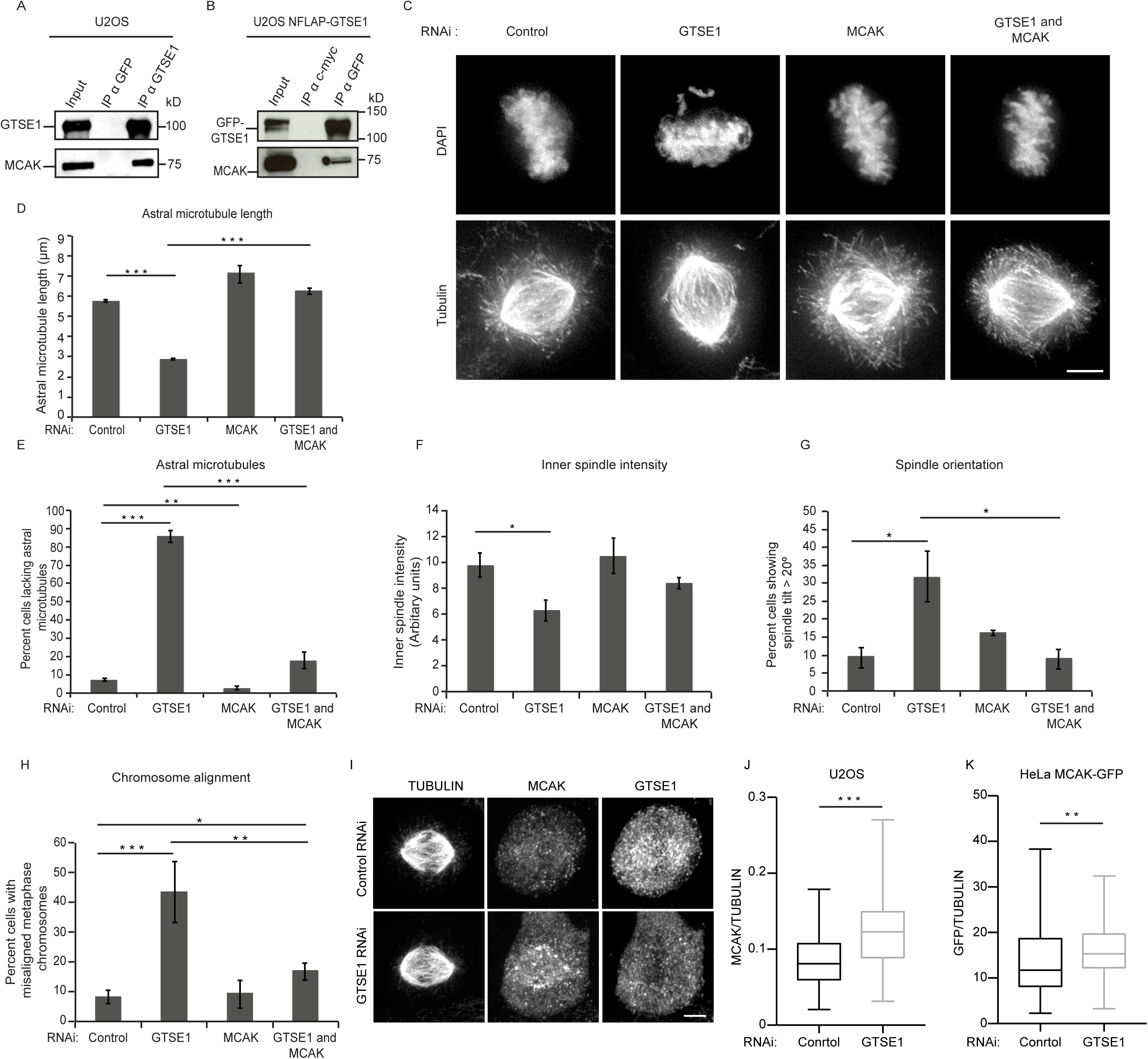
Mitotic defects following GTSE1 depletion are dependent on the activity of the MT depolymerase MCAK. (A) MCAK coimmunoprecipitates with GTSE1. Western blot following immunoprecipitation from U2OS cell lysates using either anti-GTSE1or anti-GFP antibodies, and probing with anti-GTSE1 or anti-MCAK antibodies. (B)Western blot following immunoprecipitation from U2OS NFLAP-GTSE1 cell lysates using either anti-GFP or anti-c-myc antibodies, and probing with anti-GFP or anti-MCAK antibodies. (C) Immunofluorescence images of mitotic U2OS cells followingcontrol, GTSE1, MCAK, or combined GTSE1 and MCAK RNAi, stained for DNA (DAPI) and microtubules (tubulin). (D) Quantification of the average length of astral microtubules in three dimensions from immunofluorescence analysis as shown in c.10 astral MTs were measured per cell. n = 10 cells per experiment, 3 experiments per condition. (E) Quantification of the percent cells lacking astral microtubules from immunofluorescence analysis as shown in c. n > 100 cells per experiment; 3experiments per condition. (F) Quantification of “inner-spindle” tubulin fluorescence intensity from immunofluorescence analysis as shown in C. n ≥ 19 per experiment; 3experiments per condition. (G) Analysis of spindle orientation. Quantification of thepercent of metaphase cells with a spindle tilt angle greater than 20 degrees is shown for each RNAi condition. n > 140 cells over 3 experiments per condition. (H)Quantification of the percent cells with misaligned chromosomes from immunofluorescence analysis as shown in C. n > 100 cells per experiments; 3experiments per condition. (I) Immunofluorescence images of mitotic U2OS cells showing MCAK localization after control and GTSE1 RNAi. Cells were stained for MCAK, GTSE1 and microtubules (tubulin). (J) Quantification of MCAK intensity on theinner spindle normalized to tubulin intensity in U2OS cells using immunofluorescence images after control and GTSE1 RNAi. n ≥ 105 cells over 3independent experiments. P-values were obtained using Mann-Whitney U test. (K)Quantification of MCAK intensity using GFP fluorescence on the inner spindlenormalized to tubulin intensity in HeLa cells expressing BAC based MCAK-GFP using immunofluorescence images after control and GTSE1 RNAi. n ≥ 89 cells over 3independent experiments. P-values were obtained using Mann-Whitney U test. All scale bars represent 5 micrometers. All error bars represent standard error of themean. p≤0.05 *, p≤0.01 **, p≤0.001 ***

The perturbed mitotic phenotypes following GTSE1 depletion (loss of MT stability and chromosome alignment defects) are reminiscent of the reported mitotic phenotypes upon increasing MCAK activity via overexpression (Maney et al., 1998; Moore and Wordeman, 2004; Zhang et al., 2011), suggesting that GTSE1 may normally attenuate MCAK activity, which becomes unregulated and hyperactive upon loss of GTSE1. We reasoned that if the loss of microtubule stability in mitosis following GTSE1 depletion resulted from excessive MCAK activity, then reducing MCAK levels/activity in these cells should restore MT stability. To test this, we depleted GTSE1 and MCAK by RNAi either individually or together and imaged mitotic cells by immunofluorescence (Figs. 4 C and S1, E and F). RNAi of MCAK alone in U2OS cells strongly decreased MCAK protein levels and led to an increase in mitotic cells containing longer, curved, and more dense astral MTs, consistent with previous reports (Rankin and Wordeman, 2010; Rizk et al., 2009). When MCAK was codepleted from cells with GTSE1, in contrast to GTSE1 depletion alone, most cells displayed abundant, long astral MTs, indicating that the loss microtubule stability observed after GTSE1 depletion is dependent on MCAK activity (Fig. 4 C-E). Microtubule intensity within the inner spindle regions also increased after co-depletion of GTSE1 and MCAK as compared to GTSE1 alone, indicating this was not specific to astral MTs (Fig. 4F). Because astral MTs are known to mediate spindle orientation (Pearson and Bloom, 2004), we asked whether the spindle orientation defect after GTSE1 RNAi was restored after MCAK co-depletion as well. Indeed, the spindle orientation defect following GTSE1 RNAi was also dependent on MCAK (Fig. 4 G). If the defects in chromosome alignment seen upon depletion of GTSE1 resulted from MCAK-mediated defects in microtubule stability, we expected they would also be ameliorated by co-depletion of MCAK. Remarkably, upon co-depletion of GTSE1 and MCAK, there was a dramatic reduction in misaligned chromosomes as compared to GTSE1 RNAi alone (Fig. 4H).

The above results show that all observed mitotic phenotypes associated with depletion of GTSE1 are alleviated by co-depletion of MCAK. The same effect was seen when MCAK was depleted in GTSE1^KO^ cells (Fig S2 I). This alleviation was specific to MCAK, as co-depletion of the related Kinesin-13 MT depolymerase Kif2A with GTSE1 did not restore MT stability nor chromosome alignment (Figs S3 C and S1 G). Furthermore, although total MCAK proteins levels were unchanged following loss of GTSE1 (Fig S1 H), MCAK localization to the spindle is abnormal (Fig 4I). Together, the above findings indicate that GTSE1 functions to negatively regulate MCAK microtubule depolymerase activity to ensure stability of microtubules throughout the spindle required for spindle orientation and chromosome alignment.

### GTSE1 interacts directly with MCAK and inhibits its microtubule depolymerase activity in vitro

To determine the mechanism by which GTSE1 antagonizes MCAK activity, we first asked if we could detect a direct interaction of MCAK with a defined domain of GTSE1. We performed *in vitro* pull-down assays with purified MCAK and either a purified GST-tagged GTSE1 N-terminal fragment containing residues 1-460, or a C-terminal fragment containing residues 463-739, the latter of which contains the regions of GTSE1 previously identified to interact with EB1 (Scolz et al., 2012) and p53 (Monte et al., 2003). The N-terminal fragment abundantly pulled down purified MCAK protein, while the C-terminal fragment did not (Figs. 5 A and S4 A), indicating that GTSE1 binds to MCAK via the N-terminal half of the protein.

**Figure 5.**
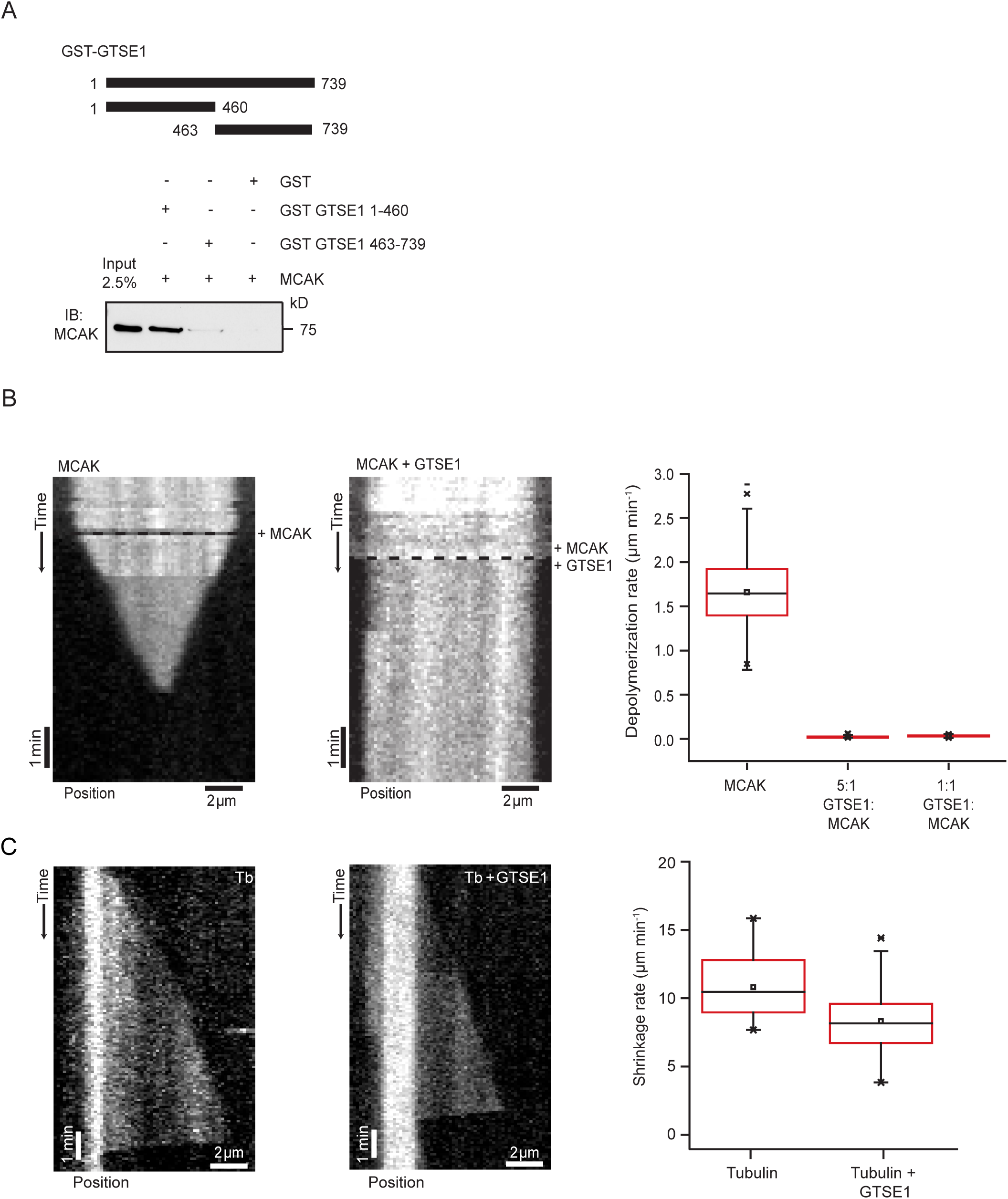
GTSE1 interacts directly with MCAK and inhibits its microtubule depolymerase activity in vitro. (A) Immunoblot showing MCAK pulled down in an in vitro GST pull down using GST alone, GST-GTSE1 1-460, and GST-GTSE1 463-739 fragments. Input represents 2.5% of total MCAK protein used for GST pull down assay. (B) Kymograph depicting 50 nM MCAK depolymerizing a GMPCPP stabilized microtubule (Supplementary movie 5). The black dashed line represents the start ofthe experiment when MCAK was added. Second kymograph depicts a GMPCPP stabilized microtubule maintaining constant length in the presence of 50 nM MCAK plus 250 nM GTSE1 (Supplementary movie 6). The black dashed line represents the start of the experiment when MCAK and GTSE1 were added. Depolymerization rates of GMPCPP stabilized microtubules in the presence of 50 nM MCAK alone, 50 nM MCAK with a 5 fold excess of GTSE1, and 50 nM MCAK with an equimolar amount ofGTSE1 are shown in the box plot. n >100 microtubules over 4 experiments. (C) Kymographs of microtubule growth and catastrophe in the presence of 20 μM tubulin alone or 20 μM tubulin and 250 nM GTSE1 from a GMPCPP microtubule seeds. Box plot shows the the shrinkage rate following catastrophe of 20 μM tubulin alone and 20 μM tubulin with 250 nM GTSE1, n = 34 and 90 microtubules, respectively.

To test whether GTSE1 could inhibit MCAK activity *in vitro*, we purified full-length GTSE1 and MCAK from insect cells (Fig S4 B-D) and assayed the MT depolymerase activity of MCAK on labeled MTs using total internal reflection fluorescence (TIRF) microscopy (Helenius et al., 2006). Addition of MCAK alone to GMPCPP-stabilized MTs resulted in rapid MT depolymerization (Figure 5B, Video 5). Addition of either equimolar or 5-fold excess amounts of GTSE1 protein to this reaction completely eliminated MCAK-induced depolymerization (Fig. 5B; Video 6). This was not due to an inherent ability of GTSE1 to stabilize MTs, because GTSE1 does not have the same impact on MCAK-independent MT depolymerization. We could only detect a minimal reduction of MT shrinkage rate following catastrophe when the higher concentration of GTSE1 alone was added to dynamic MTs, and catastrophe frequency was not reduced (Fig. 5C, data not shown). Consistently, we also observed a concentration-dependent inhibition of MCAK depolymerase activity using taxol-stabilized MTs and decreasing amounts of GTSE1 protein in bulk MT sedimentation assays (Desai et al., 1999) (Fig. S4 E). Thus, GTSE1 inhibits MCAK depolymerization activity *in vitro*.

### Depleting GTSE1 reduces chromosome missegregation frequency in CIN cancer cell lines

Overexpression of MCAK in CIN cancer cell lines reduces MT-kinetochore attachment hyperstability, thereby allowing correction of erroneous merotelic MT-kinetochore attachments and decreasing the rate of chromosome missegregation and CIN (Bakhoum et al., 2009b). Because GTSE1 is a negative regulator of MCAK activity, and has upregulated protein levels in several cancer cell lines and tumours relative to non-transformed cells (Scolz et al., 2012), we wondered whether GTSE1 expression in these cells contributed to CIN. First, we analyzed anaphase chromosome segregation defects, including lagging chromosomes, which correlate with CIN (Thompson and Compton, 2008; Bakhoum et al., 2014). In HeLa and U2OS cells, highly CIN cancer cell lines with high levels of GTSE1 protein, we found that reduction of GTSE1 levels by RNAi significantly reduced the frequency of anaphase chromosome segregation defects (Fig. 6 A-C). Remarkably, both U2OS cell lines with GTSE1 stably knocked out showed an even greater reduction in the frequency of anaphase chromosome segregation defects (Fig. 6 C). Closer analysis of GTSE1-depleted cells revealed that the reduction in anaphase chromosome segregation defects could be attributed specifically to lagging chromosomes, which generally arise from merotelic attachments, as opposed to chromosome bridges or acentric fragments (Fig. S 5). Importantly, this impact of GTSE1 on chromosome missegregation is also mediated through MCAK, as reducing GTSE1 levels in cells also depleted of MCAK did not have an affect on the frequency of anaphase chromosome segregation defects (Fig. 6 C).

**Figure 6.**
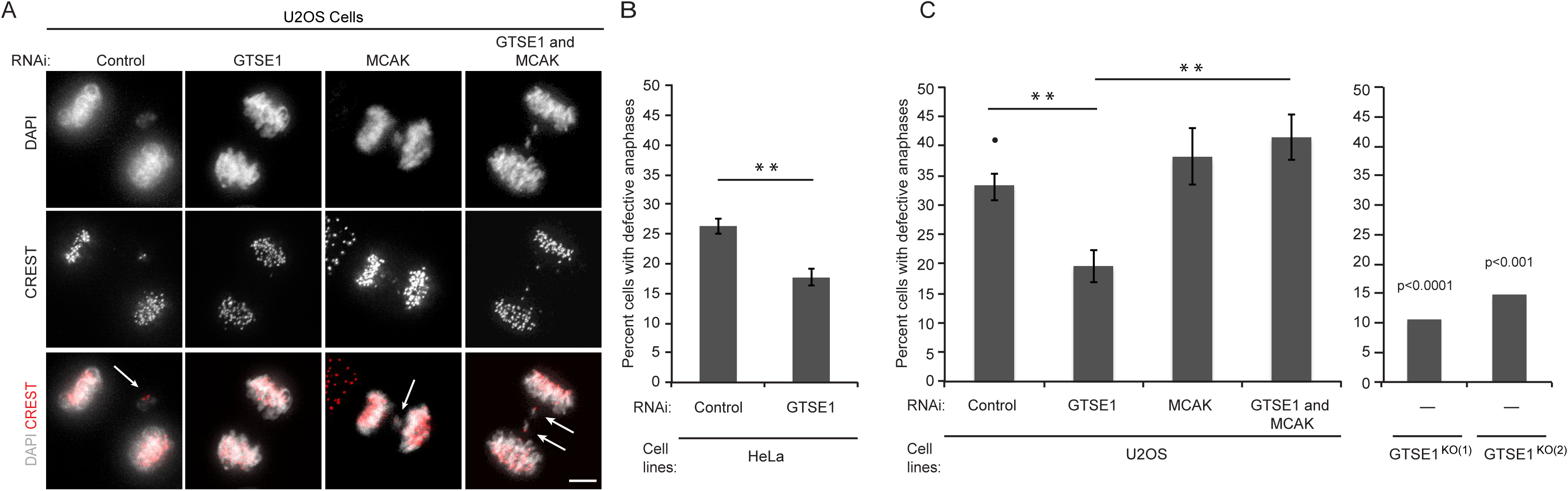
Reduction of GTSE1 levels reduces anaphase chromosome segregation defects in HeLa and U2OS cells. (A) Immunofluorescence images of anaphase U2OS cells after control, GTSE1, MCAK, or GTSE1 and MCAK siRNA, stained for DNA (DAPI) and kinetochores (CREST). (B) Quantification of the percentage of anaphase HeLa cells with defective anaphase chromosome segregation events. n > 390 per condition over 3 experiments. (C) Quantification of the percentage of anaphase U2OS cells with defective anaphase chromosome segregation events from immunofluorescence analysis as shown in A.. The left bar chart shows U2OS cells after RNAi conditions as in A. n > 65 per experiment; 3 experiments per condition. The right bar chart shows two independent U2OS GTSE1 knockout clones. P-values were obtained from a chi-squared test comparing to the U2OS control condition (marked with •). n > 100 cells.p≤0.05 *, p≤0.01 **, p≤0.001 ***

### Elevating GTSE1 levels induces chromosome missegregation and chromosomal instability

We next asked if overexpression of GTSE1 induced anaphase chromosome segregation defects in HCT116 cells, which are near-diploid, relatively chromosomally stable, and have relatively low levels of GTSE1 protein (Fig. 7 A) (Thompson and Compton, 2008). We stably transfected HCT116 and HCT116 p53 ^-/-^ cells with a GTSE1-GFP cDNA construct that allowed us to maintain cells overexpressing GTSE1 over many generations through antibiotic selection (Fig. 7 A, B). We isolated clonal transformants overexpressing GTSE1-GFP approximately 3 to 10-fold higher than non-transfected cells and grew them for approximately 60 generations before analyzing mitotic cells. Although these cells displayed an approximate 30% increase in MT density, no major morphological changes to the spindle were observed and relative TACC3 recruitment to MTs did not increase (Fig S3 D). All HCT116 cell clones overexpressing GTSE1 did, however, show a significant increase in the frequency of anaphase chromosome segregation defects (Fig. 7, C and D).

**Figure 7.**
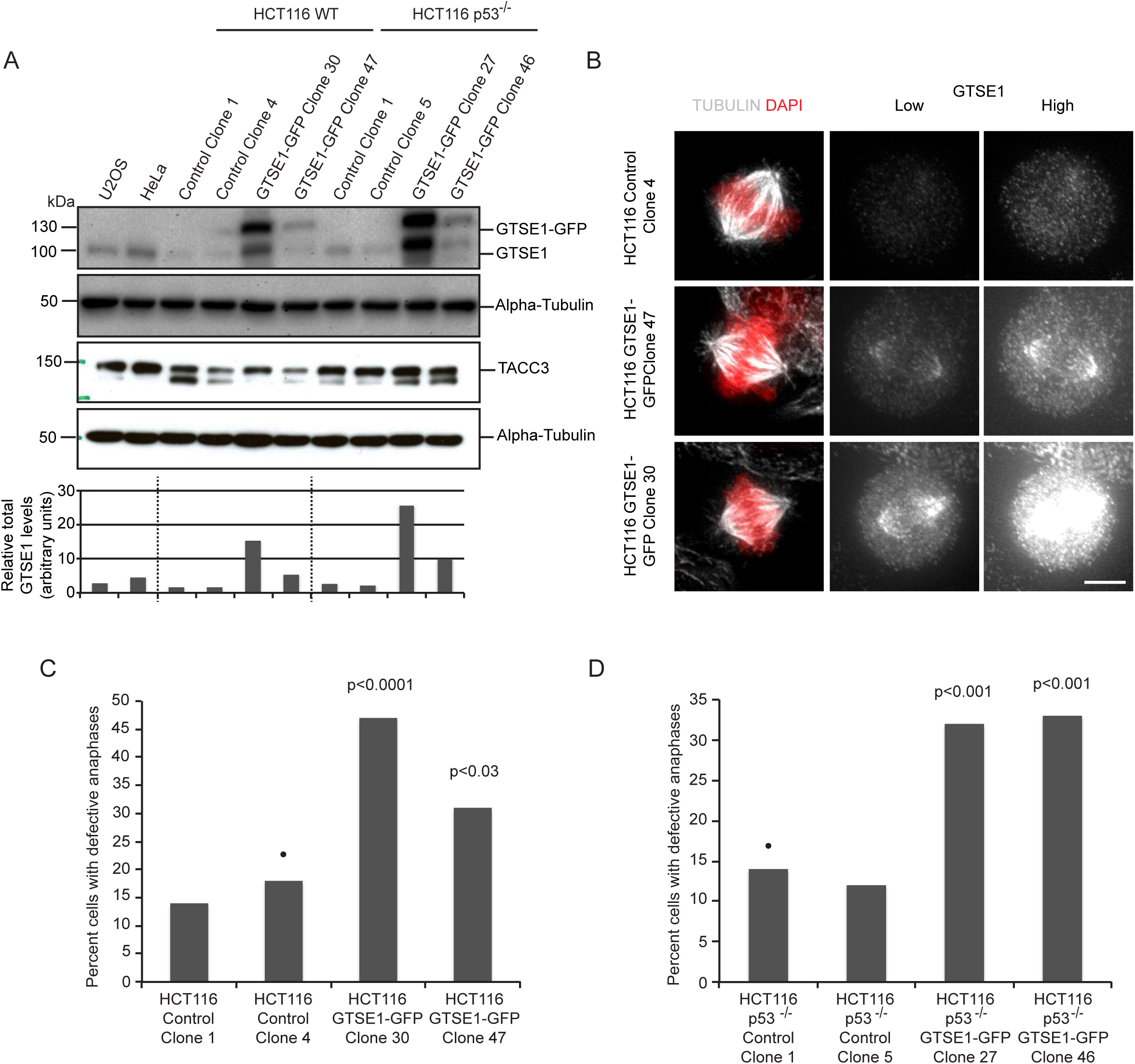
Overexpression of GTSE1 induces segregation defects in HCT116 cells.(A)Representative Western blots of cell lysates from U2OS, HeLa, and HCT116 and HCT116 p53-/-control and GFP-GTSE1 expressing clonal cell lines used for analysis in (C) and (D). Blots were probed with anti-GTSE1, anti-TACC3 and anti-alpha-tubulin. The box chart represents quantification of GTSE1 protein levels normalized to tubulin levels from the blot shown. The relative abundance of total GTSE1 protein was quantified and normalized to tubulin levels. (B) Low and high intensity immunofluorescence images of HCT116 control and GTSE1-GFP overexpressing clones showing normal spindle morphology stained for DNA (DAPI), GTSE1 and microtubules (tubulin). (C) Quantification of the percent of anaphase cells with lagging chromosomes for control or GTSE1-GFP expressing HCT116 clones. n ≥ 99; P-values were obtained from chi-squared tests comparing control clones designated with •. (D) Quantification of the percent of anaphase cells with lagging chromosomes for control or GTSE1-GFP expressing HCT116 p53-/-clones. n ≥ 104; P-values were obtained from chi-squared tests comparing control clones designated with •. p≤0.05*, p≤0.01 **, p≤0.001 ***

Finally, we asked whether increased levels of GTSE1 in HCT116 cells would induce CIN in these cells. Clonal cell lines overexpressing GTSE1 were analyzed for the frequency with which cells contained deviations from the disomic state after approximately 60 generations by fluorescence in situ hybridization (FISH) analysis of chromosomes 7 and 11 (Fig. 8 A), as compared to mock-transfected clones. Independent HCT116 clones overexpressing GTSE1-GFP displayed a significant increase in cells containing deviations from the disomic state (Figure 8 B-D). As p53 has been reported to induce apoptosis following chromosome missegregation (Thompson and Compton, 2008), we also performed these experiments in HCT116 p53 ^-/-^cells, which also showed elevated frequencies of anaphase segregation defects after GTSE1 overexpression (Fig. 7 D). HCT116 p53-/-clones overexpressing GTSE1-GFP displayed a yet greater increase in deviations from the modal number of chromosomes as compared to mock-transfected clones (Fig. 8B, E, F). Thus, reduction of GTSE1 levels in highly CIN cancer cell lines reduces the frequency of lagging chromosomes, while overexpression of GTSE1 in chromosomally stable cell lines induces lagging chromosomes and whole chromosome CIN.

**Figure 8.**
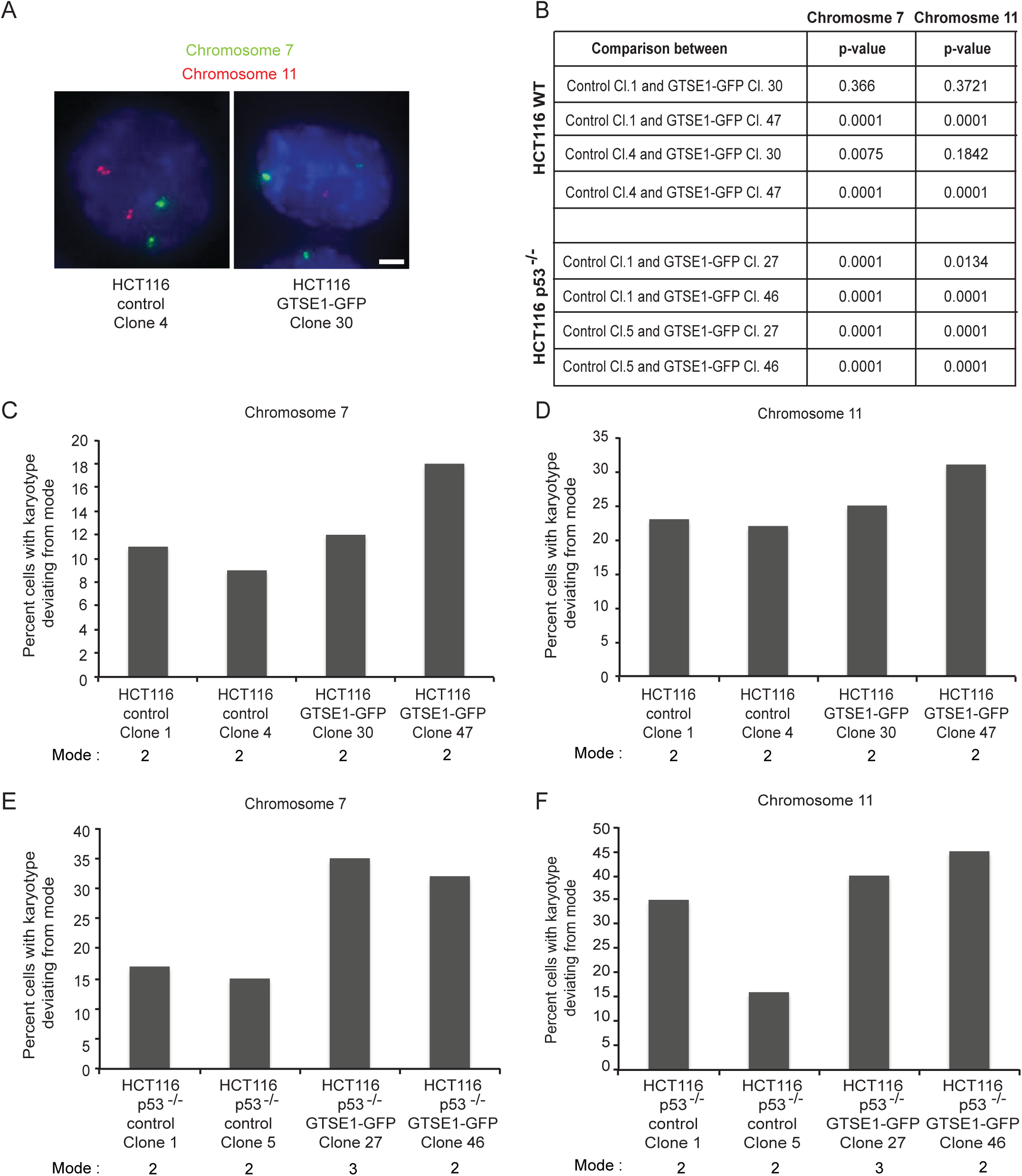
Overexpression of GTSE1 induces CIN in HCT116 cells. (A) Fluorescence images from fixed control or GTSE1-GFP overexpressing clonal HCT116 cells processed forfluorescence in situ hybridization (FISH). Probes for α-satellite regions on Chromosomes 7(green) and 11 (red) are shown and were used to count the number of copies of each chromosome in cells. (B) Statistical significance values from comparing deviances from themodal number of chromosomes in control versus GTSE1-GFP overexpressing HCT116 cellsfrom data presented in (C-F). P-values were determined from chi-squared analysis comparing the indicated clonal cell lines. (C) Percentage of HCT116 control or GFP-GTSE1 expressing clonal cell lines containing numbers of chromosome 7 deviating from the mode. n > 980 cells for each condition. (D) Percentage of HCT116 control or GFP-GTSE1 expressing clonal cell lines containing numbers of chromosome 11 deviating from the mode. n > 980cells for each condition. (E) Percentage of HCT116 p53-/-control or GFP-GTSE1 expressing clonal cell lines containing numbers of chromosome 7 deviating from the mode. n > 1000 cells for each condition. (F) Percentage of HCT116 p53-/-control or GFP-GTSE1 expressing clonal cell lines containing numbers of chromosome 11 deviating from the mode. n > 1000 cells for each condition.

## Discussion

Here we have demonstrated that GTSE1 regulates MT stability by inhibiting MCAK MT depolymerase activity. Depletion of GTSE1 enhances MCAK activity in mitotic cells to levels that are incompatible with the stabilization of astral microtubules and MT-kinetochore interactions, leading to defects in spindle positioning and chromosome capture and alignment. At the same time, GTSE1 depletion results in less chromosome segregation defects in the cancer cell lines tested, consistent with the decreased stability of kinetochore-MT interactions enhancing the ability to correct merotelic attachments. Remarkably, knocking out GTSE1 in U2OS cancer cells resulted in stable cell lines with dramatically reduced missegregation frequencies. While these knockout cell lines also displayed decreased MT stability similar to that observed after RNAi-mediated depletion of GTSE1, they did not exactly phenocopy siRNA-depleted cells: in knockout cells chromosome misalignment was less severe. Because the BAC transgene rescue experiments (Figures 1 B and 2 D) ruled out off-target siRNA effects, it appears that this difference in the knockout cells reflect either a long-term GTSE1-loss phenotype, or that clones have been “adapted” though selection to tolerate loss of GTSE1 by suppressing the chromosome misalignment defect. Of note, we did not observe reduction in MCAK protein levels in these cells, and the dependency on MCAK of the perturbed phenotypes remained.

The precise regulation of MCAK activity is essential for a number of cellular processes that rely on dynamic microtubules (Maney et al., 1998; Rankin and Wordeman, 2010) (Kline-Smith et al., 2004; Lan et al., 2004; Braun et al., 2014; Walczak et al., 2002; Domnitz et al., 2012). The potent MT depolymerase activity of MCAK observed *in vitro* suggests that its activity must be generally repressed in cells. This may allow for precise spatial and temporal activation of its activity for discrete functions. Here we have elucidated a novel regulatory mechanism responsible for inhibition of MCAK activity, through interaction with GTSE1. *In vitro*, GTSE1 protein present at equimolar amounts to MCAK is sufficient to completely inhibit MCAK depolymerase activity, suggesting GTSE1 may be a target for differential regulation of MCAK activity in cells.

Most proposed mechanisms to date for direct inhibition of MCAK activity involve phosphorylation of MCAK and/or intramolecular rearrangements (Ems-McClung et al., 2013; Talapatra et al., 2015; Burns et al., 2014). Several phosphorylation sites dependent on multiple mitotic kinases identified on MCAK have been shown to be required for both positive and negative modulation of its activity (Lan et al., 2004; Zhang et al., 2011; Tanenbaum et al., 2011; Ems-McClung et al., 2013; Andrews et al., 2004; Sanhaji et al., 2010; Zhang et al., 2007), but how these various phosphorylation events control MCAK activity in the cellular context remain largely unknown. It was recently shown that NuSAP modulation of MCAK activity is dependent on Aurora B phosphorylation (Li et al., 2015). It will be interesting in the future to determine if GTSE1 association with MCAK is regulated by, or influences, MCAK’s phosphorylation state. A conformation of MCAK where the C-terminal tail interacts with its motor/neck domain is emerging as an important transition state during its catalytic cycle, and this transition could be a target for controlling its cellular regulation (Ems-McClung et al., 2013; Talapatra et al., 2015; Zong et al., 2016). Indeed, phosphorylation by Aurora B has been shown to affect this conformation in vitro (Ems-McClung et al., 2013). Furthermore, it was recently shown that point mutations in MCAK impairing this conformation lead to increased MCAK activity toward MTs, due to enhanced accumulation of MCAK at the spindle poles, reminiscent of what we observe following GTSE1 depletion (Zong et al., 2016) (Fig. 3 I). Determining the impact of GTSE1 interaction with MCAK on its ability to transition between distinct conformational states will be informative.

Our finding that GTSE1 regulates MT stability is interesting to consider in the context of Aurora A kinase control of MT stability. Aurora A kinase activity controls GTSE1 localization to the spindle (Hubner et al., 2010). During mitosis, Aurora A phosphorylates the TACC3 protein, which allows TACC3 to interact with clathrin heavy chain (CLTC), forming a microtubule-interaction interface that brings the pTACC3-clathrin complex to the spindle (Hubner et al., 2010; Booth et al., 2011; Hood et al., 2013; Lin et al., 2010; Fu et al., 2010; Kinoshita, 2005). GTSE1 spindle localization is dependent on Aurora A, TACC3, and clathrin, and GTSE1 is the most downstream of these components recruited to the spindle (Hubner et al., 2010). Loss of Aurora A, TACC3, or clathrin, like GTSE1, results in defects in MT stability and chromosome alignment (Lin et al., 2010; Fu et al., 2010; Booth et al., 2011; Marumoto et al., 2003; Bird and Hyman, 2008; Giet, 2002). The TACC3-clathrin complex has been shown to be required for kinetochore MT integrity and the presence of a “mesh” of inter-MT connections within kinetochore MT bundles (Booth et al., 2011; Hood et al., 2013; Nixon et al., 2015). This “mesh” has been proposed to also include GTSE1 and serve to physically stabilize MTs within these bundles (Nixon et al., 2015). Our results suggest that Aurora A-dependent recruitment of the TACC3-clathrin complex also facilitates MT stability by recruiting GTSE1 to inhibit MCAK.

While hyperstabilization of kinetochore-microtubule dynamics has been shown to influence chromosomal instability in cancer cells, the genetic and molecular defects which contribute to MT hyperstabilization and CIN in tumours remain unclear. Sequencing of cancer genomes has revealed that mutations in genes directly controlling chromosome segregation and mitosis are relatively rare (Orr and Compton, 2013). Rather, it appears that CIN may originate from changes in protein expression or activity due to perturbed oncogenic signaling pathways (Duijf and Benezra, 2013; Orr and Compton, 2013). We show here that changing GTSE1 expression levels influences chromosome segregation and CIN. Deregulation of GTSE1 expression levels could therefore represent a molecular defect contributing to the stabilization of MT dynamics and induction of CIN in tumours through downregulation of MCAK activity. Induction of CIN in tumour cells is likely the result of complex changes in oncogenic pathways and coordinated gene expression, however, necessitating care in relating single gene perturbations in model systems to the tumour condition (Thiru et al., 2014). Our work also predicts that GTSE1 could be a CIN–related effector of increased Aurora A activity. Aurora A and its activator TPX2 are commonly found overexpressed in tumours, highly associated with CIN (Carter et al., 2006), and proposed to comprise a oncogenic holoenzyme (Asteriti et al., 2010). Overexpression of Aurora A has recently been shown to induce kinetochore MT hyperstability and CIN in colorectal cancer cells, and these affects were attributed to increased MT polymerization rates (Ertych et al., 2014). Because GTSE1 spindle recruitment is dependent on Aurora A activity, it is possible that overexpression of Aurora A also results in enhanced recruitment of GTSE1 to the spindle and increased inhibition of MCAK, which in turn could alter MT dynamics and stabilize kinetochore MTs, promoting CIN.

We have described here a novel role of GTSE1 in regulating chromosome stability through microtubule stability. Inhibition of MCAK activity by GTSE1 provides a new mechanism by which cells tune MT dynamics to ensure the precise balance of MT stability required for chromosome alignment and segregation.

## Materials and methods

### Cloning and Plasmids

The hGTSE1-GFP-T2A-BSD plasmid was generated by in-frame cloning of *GTSE1* cDNA, eGFP and T2A-BSD in a EGFP-N2 vector (Clontech). The GST-hGTSE1 construct used for protein purification from insect cells was generated by cloning *GTSE1* cDNA using EcoRI and BamHI restriction sites into pFLMultiBac vector modified by adding GST (Van Stedum and King, 2002; Fitzgerald et al., 2006). The GST-hGTSE1 fragment expression constructs were generated by sub-cloning the sequences encoding amino acids 1-460 (N-terminus) and 463-739 (C-terminus) in a pGEX-6p-1rbs vector (GE Healthcare) using BamHI and SalI restriction sites. An N-terminal FLAG-LAP “NFLAP” cassette was recombined onto the N-terminus of *GTSE1* encoded on the BAC RP11-1152E11 via Red E/T-based recombination (Poser et al., 2008; Zhang et al., 1998). An mCherry-BSD cassette was recombined onto the C-terminus of mH2A.Z encoded on the BAC RP24-363J17 via Red E/T-based recombination (Poser et al., 2008; Zhang et al., 1998).

### Cell culture and cell lines

All cell lines were grown in DMEM (PAN Biotech) containing 10% fetal bovine serum (Gibco), 2 mM L-Glutamine (PAN Biotech), 100U/mL Penicillin and 0.1mg/ml streptomycin (PAN Biotech) at 37 °C in 5% CO2. U2OS cells stably expressing RNAi-resistant wild-type and SxIP-mutated GTSE1-LAP were described previously(Scolz et al., 2012). HCT116 p53-/- and U2OS-PA-GFP-tubulin cells were kind gifts from Duane Compton. U2OS cells expressing NFLAP-GTSE1, GTSE1-LAP^WT(303)^, and mH2A.Z-mCherry were generated by transfecting the respective BACs using Effectene according to manufacturers protocol, and selecting for stable transfectants. HeLa cells expressing bacterial artificial chromosome based hKif2C-LAP were a kind gift from Anthony Hyman. HCT116 and HCT116 p53^-/-^ cells grown in 6 cm dishes were transfected with 1.5 μg of hGTSE1-GFP-T2A-BSD cDNA using 5 μL of Lipofectamine 2000 (Invitrogen) according to manufacturer’s protocol. Stable line populations were selected on BSD (4 μg/mL), and individual clones isolated. HCT116 and HCT116 p53-/-control clones were isolated after treating the cells with Lipofectamine 2000 in absence of DNA.

### Generation of GTSE1 knockouts using CRISPR-Cas9 system

To generate *GTSE1* knockout cells using the CRISPR-Cas9 system, two different target sites were chosen in *GTSE1* downstream of the ATG in exon 2(GGCAGGCTGAAGGCTCATCG) and exon 9 (AAGGCCAGAGCAGCGCCGGC). Plasmids expressing the Cas9 nuclease and the corresponding guide RNAs were obtained by cloning the following primers pairs: (5’-CACCGAAGGCCAGAGCAGCGCCGGC-3’ and 5’-AAACGCCGGCGCTGCTCTGGCCTTC-3’) and (5’-CACCGGCAGGCTGAAGGCTCATCG-3’ and 5’-AAACCGATGAGCCTTCAGCCTGCC-3’)into the pX330 background (from Addgene), as described in(Ran et al., 2013). U2OS cells were transfected with individual plasmids targeting a single site, or with two plasmids targeting both sites simultaneously, and clones were isolated by serial dilution 48 h after transfection. Genomic DNA was prepared using the DNeasy Blood and Tissue kit (Qiagen). The regions surrounding the CRISPR-Cas9 sites were amplified using the primers 5’-CGGCAATGAGTCTCCCTCAG-3’ and 5’-AATCGCTTGAACCCGAAAGG-3’ (for the site in exon 2) or 5’-CAGTCCACAGCAAATGCCAG-3’ and 5’-CACGACTGAGGTGTGACTTC-3’ (for the site in exon 9) and the corresponding PCR products were sequenced using primers 5’-CCCTGGGATGCGATCATTTC-3’ or 5’-CCCTCAGCACTGCATTAGCAC-3’, respectively. Heterozygous insertions and deletions were resolved using the TIDE software (http://tide.nki.nl) (Brinkman et al., 2014). TIDE compares sequence traces (from capillary sequencing runs) between a reference run (wild type sequence) and a sample run (modified sequence) to identify mutations, insertions and deletions in the sample sequence. It also calculates the proportion of modifications at a given site in the modified DNA.

### Generation of GFP knock in at endogenous GTSE1 via Cas9/CRISPR

To generate the U2OS GTSE1-GFP-knock in, a BAC containing a GTSE1-GFP-T2A-BSD^R^ fusion was first generated via Red E/T-based recombination of a GFP-T2A-BSD^R^ cassette (gift of Tony Hyman) amplified with the primers TGAGCCCTGAGGCTGACAAGGAGAACGTGGATTCCCCACTCCTCAAGTTCGAGAATCTTTATTTTCAGGGCG and CAAGTGTAAGCCACTGCGACCAGCCTAAAGCTTGTTTCTGAGGGTTGAAAATTAACCCTCCCACACGTAGCC into a BAC containing GTSE1. The template for Cas9 mediated HR was obtained by amplifying from the GTSE1-GFP-T2A-BSD^R^ BAC a region encompassing the GFP-T2A-BSD^R^ cassette and 1kb phomology arms to the C-terminus of GTSE1 using primers CATCAGCCAAGTGAACGAGC and GAATCACGCAGTAACCGCAG. The plasmid expressing the Cas9 nuclease and the guide RNA targeting the C-terminus of GTSE1 was obtained by cloning the following primers pairs: (CACCGTTGAAAGAACAGCCCTAAAG and AAACCTTTAGGGCTGTTCTTTCAAC) into the pX330 background.

### RNAi

siRNA against hGTSE1 (5′-GAUUCAUACAGGAGUCAAA-3′), hMCAK (5′-GAUCCAACGCAGUAAUGGU-3′) and control siRNA (Silencer negative control #2; product number AM4637) were purchased from Ambion. siRNA against hKif2A (#2 5’-CUACACAACUUGAAGCUAU-3’, #4 5’-GACCCUCCUUCAAGAGAUA-3’) were purchased from Sigma-Aldrich. Approximately 35,000 U2OS cells were added to prewarmed media in 24 well plates or 8-well imaging chambers (ibidi), and transfection complexes containing 2.5 μl oligofectamine and siRNA were added immediately afterwards. Media was changed after 6–8 h. A final concentration of 80 nM and 12 nM RNAi were used for GTSE1 and MCAK, respectively. For Kif2A, a final concentration of 100 nM was used by mixing siRNA #2 and #4. All experiments were performed 48 hours after siRNA transfection.

### Antibodies

Rabbit antibodies against hGTSE1 were previously described (Scolz et al., 2012). Goat anti-GFP antibodies used in immunoprecipitation assays were previously described (Poser et al., 2008). Rabbit antibody against CEP135 was previously described (Bird and Hyman, 2008). Mouse monoclonal antibody conjugated with 488 against Mad1 was a kind gift from Andrea Musacchio. Rabbit anti-pTACC3 was a kind gift from Kazu Kinoshita. The following antibodies were obtained from commercial sources: mouse anti-alpha-tubulin (DM1alpha, Sigma Aldrich), mouse anti-MCAK (Abnova Corporation, Clone 1G2), human nuclear antibodies to nuclear antigens–centromere autoantibody (CREST; CS1058; Europa Bioproducts Ltd.), mouse anti-c-myc (Oncogene/Calibiochem), rat anti-EB1 (Absea Biotechnology, Clone KT-51), rabbit anti-TACC3 (Santa Cruz Biotechnlogy, H-300), rabbit antich-TOG (Abcam #86073), rabbit anti-Aurora B (Abcam #2254), rabbit anti-Kif2A (Novus Biologicas NB500-180), and rabbit anti-GFP (Abcam #AB6556). The following Secondary antibodies were used: donkey anti–mouse, -rabbit, or -rat conjugated to Alexa 488, 594, or 647 (Bethyl laboratories) and donkey anti–human conjugated to Texas Red or Cy5 (Jackson ImmunoResearch Laboratories).

### Immunofluorescence

Cells on coverslips were fixed using −20 °C methanol for 10 minutes. Cells were blocked with 0.2% fish skin gelatin (Sigma-Aldrich) in PBS. Cells were incubated with primary antibodies in 0.2% fish skin gelatin in PBS for 1 hour at 37 °C in a humidified chamber, washed, and the same repeated with secondary antibodies. Coverslips were mounted with ProLong gold with DAPI (Molecular Probes, Life technology). Quantification of TACC3 on inner spindle of U2OS cells after control and GTSE1 RNAi and in GTSE1 knockout clones was performed following fixation in 4% PFA dissolved in PIPES (50 mM PIPES pH 7.2m 10 mM EGTA, 1 mM MgCl2, 0.2 % Triton X-100.

### Microscopy and live cell imaging

Fluorescence dissipation after photoactivation (FDAPA) analysis and images used for quantifying astral length, inner spindle intensity, spindle orientation, and TACC3 and MCAK spindle localization were acquired using a Marianas (3i)spinning disk confocal system based on an Axio Observer Z1 microscope (Zeiss) equipped with a Hamamatsu ORCA-Flash 4.0 Camera. Images were taken using 63× 1.4 NA Apochromat objective (Zeiss). Images for TACC spindle localization in U2OS cells were obtained using a 100× 1.4 NA Plan-Apochromat objective (Zeiss). The images were Z-projected using Slidebook software 5.5. All other images were acquired using a DeltaVision imaging system (GE Healthcare) equipped with an sCMOS camera (PCO edge 5.5). Images were taken using a 60× 1.42 NA PlanApo-N objective (Olympus) at room temperature. Serial Z-stacks of 0.2 μm thickness were obtained and deconvolved using SoftWoRx 6.1.1 software. For live cell imaging, media was changed to CO2 independent media (Gibco) 12 hours prior to imaging. Live cell image sequences were acquired at 1 min intervals for 12 hours in 2 μm serial Z sections using a 40× 1.42 NA UPlanFL-N objective (Olympus) at 37 °C.

### Image quantification and data analysis

Inner spindle intensity and TACC3 and MCAK spindle localization were quantified in three dimensions using the surface module in IMARIS software (Bitplane). Astral microtubule lengths were measured in three dimensions using IMARIS software. Astral microtubule length measurements using EB1 comets were performed as described in (Stout et al., 2011). Briefly, the position of every EB1 comet was determined using the point function of the IMARIS software, and the angle it formed with the spindle axis was calculated. Each comet with an angle superior to the one formed by the spindle axis and the outermost spindle microtubule was considered as an astral comet, and its distance to the closest pole was calculated. To measure spindle tilt, both spindle poles were located in the Z-series and then using the angle tool in Slidebook software 5.5, the angle made by the spindle to the substratum was measured. Images were processed with ImageJ or Photoshop (Adobe). Total number of EB1 astral microtubule comets were quantified using IMARIS software. To measure spindle tilt, both spindle poles were located in the Z-series and then using the angle tool in Slidebook software 5.5, the angle made by the spindle to the substratum was measured. Images were processed with ImageJ or Photoshop (Adobe).

### Determining kinetochore-MT half life by FDAPA

FDAPA in U2OS cells expressing Photoactivatable (PA)-GFP-tubulin was performed as described in (Bakhoum et al., 2009b). U2OS-PA-GFP-Tubulin cells grown on poly-L-Lysine (Sigma-Aldrich) coated 3.5 cm glass bottom chamber (ibidi) were treated with either control or GTSE1 RNAi for 48 h. The medium was changed to CO2-independent media 12 h prior to the experiment. PA-GFP-tubulin in a small area around the kinetochore-MT attachment region on the metaphase spindle was activated with a 405 nm laser and images were taken every 10 sec for 5 minutes. Fluorescence intensity after activation was measured at each time point using ImageJ. Fluorescence intensity was measured within a similar region on the other non-activated half spindle was used for background subtraction. Values were corrected for photobleaching by normalizing to values obtained from taxol-treated stabilized spindles. Following background subtraction and correction for Photobleaching, values were normalized to the fid were fitted to a double exponential decay curve *F* = *A1* × exp(−*k*1 × *t*) + *A2* × exp(−*k*2 × *t*), using Prism, where A1 and A2 are the percent total fluorescence contribution of the non-kinetochore and kinetochore microtubules, k1 and k2 are their respective decay rate constants and t is the time after photoactivation. The half life of kinetochore microtubules was calculated using *T*1/2 = In 2/*k_2_*.

### Kinetochore MT stability assays and determining Mad1 positive kinetochores

U2OS cells treated with either control or GTSE1 RNAi were arrested using Cdk1 inhibitor (RO-3306, Calibiochem) at 31 hours after RNAi transfection. Cells were arrested for 17 hours and then released in normal media for 1 h. After 1 h of release the cells were immediately transferred to ice cold media for 17 minutes and fixed in −20 °C methanol. To determine the ratio of Mad1 positive kinetochores, U2OS cells treated with either control or GTSE1 RNAi were arrested using Cdk1 inhibitor (RO-3306, Calibiochem) as mentioned for kinetochore MT stability assay. After 1 hr of release the cells were fixed with 4% PFA and permeabilized with 0.1% triton X-100. The ratio of Mad1 positive kinetochores was determined by quantifying the total number of kinetochores in aligned metaphase plates and then the number of Mad1 positive kinetochores were quantified using Coloc tool of IMARIS software.

### K-Fiber localization of GTSE1

U2OS cells expressing BAC based GTSE1-LAP^WT(303)^ were arrested using Cdk1 inhibitor (RO-3306, Calibiochem) for 17 hours and then released in normal media for 1 h. After 1 h of release the cells were immediately transferred to ice cold media for 17 minutes and fixed using 4% PFA dissolved in PIPES (50 mM PIPES pH 7.2m 10 mM EGTA, 1 mM MgCl2, 0.2 % Triton X-100. GTSE1 localization to kinetochore fibers was studied using a Marianas (3i) spinning disk confocal system based on an Axio Observer Z1 microscope (Zeiss). Images were taken using a 100× 1.4 NA Plan-Apochromat objective (Zeiss).

### Immunoprecipitation

Cells with ~ 70% confluency were arrested in mitosis by adding 200 ng/mL nocodazole for 18 h. Mitotic cells were harvested by shakeoff and lysed using cell lysis buffer (50 mM HEPSES pH7.2, 50 mM Na2HPO4, 150 mM NaCl, 10% glycerol, 1% Triton X-100, 1 mM EGTA, 1.5 mM MgCl2, protease inhibitors) followed by centrifugation at 13,000 rpm for 10 minutes at 4 °C to clear the lysate. A part of the supernatant was taken as “input” and 1-2 μg of the indicated antibody was added to the remaining supernatant and incubated for 2 h at 4 °C with rotation. Dynabeads couple to protein G (Novex Life Technology) were added to the extracts and incubated for 4 hours at 4 °C. The beads were washed three times with cell lysis buffer and once with 1× PBS. The beads were resuspended in hot Lamelli buffer and analyzed by Western blotting.

### Western Blot

For western blotting after RNAi, cells were harvested by directly adding hot Lamelli buffer in 24 well plates. The proteins were separated on SDS-page gels and transferred onto nitrocellulose membranes. The membrane was incubated with the indicted primary antibodies. The secondary antibodies were coupled to horseradish peroxidase and protein bands were detected using enhanced chemiluminescence (ECL, Amersham, GE Healthcare).

### Protein Purification

Full-length human *GTSE1* cDNA sequence was cloned into pFLMultiBac vectors and baculovirus generated. hGTSE1-FL protein was expressed in Tnao38 insect cells at 27 °C for 48 hours. The cells were harvested by centrifugation at 1800 rpm for 15 minutes in a Sorvall RC 3BP+ (Thermo scientific) centrifuge. The cell pellet was either stored at −80 °C or processed immediately. Cell pellet from 1 L culture was resuspended in 100 mL ice cold Buffer A (50 mM HEPES pH 8.0, 300 mM NaCl, 5% Glycerol, 2 mM TCEP and Protease inhibitors [Serva]) lysed by sonication and clarified by centrifuging at 29,000 rpm for 50 minutes at 4 °C. The cell lysate was incubated with 1 mL Glutathione resin (Amintra) for 1 hour at 4 °C. The resin beads were loaded on gravity flow columns and washed with 150 mL Buffer A at 4 °C. hGTSE1 was cleaved from the beads using GST Precision overnight at 4 °C. The protein was eluted and concentrated using Amicon concentrators. The protein was further purified by size exclusion in a Superdex 200 10/300 column (GE Healthcare) using gel filtration buffer (30 mM HEPES pH8, 300 mM NaCl, 5% glycerol, 2 mM TCEP). The peak fractions were collected and concentrated in Amicon concentrators to give a final concentration of 5–10 μM. The hGTSE1 1-460 and 463-739 fragments and GST were cloned into pGEX-6p-1rbs vector and expressed in bacteria. Bacteria were grown to O.D. 600 of 0.8 and were induced using 1 mM IPTG at 20 °C overnight. The bacterial cells were pelleted at 4000 rpm for 20 minutes at room temperature. Cells were resuspended in GST binding buffer (25 mM HEPES pH 7.5, 300 mM NaCl, 1 mM EDTA, 5% glycerol, 1% Triton X-100, DNase), lysed by sonication and cleared by centrifugation at 29,000 rpm for 30 minutes at 4 °C. The cleared lysate was incubated with Glutathione resin (Amintra) overnight at 4 °C. The beads were washed with GST binding buffer and the protein was used for experiments. MCAK-His6 was expressed in Sf9 cells and purified by a sequence of cation exchange, nickel-affinity, and size-exclusion chromatography(Helenius et al., 2006).

### Pulldowns

*In vitro* GST pull downs were performed in GST binding buffer by incubating equal amount of purified FL-MCAK with GST alone or GST-hGTSE1 1-460 and 463-739 fragments (immobilized on Glutathione resin) for 1 h at 4 °C. The reactions were washed with GST binding buffer and resuspended in hot Lamelli buffer and analyzed by Western blotting.

### Microtubule Pelleting Assays

Taxol stabilized microtubules were prepared by incubating microtubules with 1 mM GTP at 37 °C for 30 minutes followed by incubation at 37 °C for 5 minutes after addition of 50 μM taxol. 1.66 μM tubulin was added to a reaction mixture of MCAK (200 nM) and increasing amounts of GTSE1. All reactions were performed in BRB80 buffer (80 mM Pipes pH 6.8, 1 mM EGTA, 1mM MgCl2) supplemented with 70 mM KCl, 1.5 mM ATP, 10 μM taxol for 1 h at room temperature. The reaction was then layered onto a cushion buffer (80 mM PIPES, 1 mM EGTA, 1 mM MgCl2, 50% glycerol) in a microcentrifuge tube and was centrifuged at 90,000 rpm in a TLA-120.1 rotor (Beckman-Coulter) for 10 min at 25 °C. Supernatant and pellet fractions were separated by SDS–polyacrylamide gel electrophoresis and stained with Coomassie blue for analysis.

### Total internal reflection fluorescence microscopy assays for microtubule dynamics and MCAK activity

#### Tubulin and microtubule preparation

Tubulin was purified from juvenile bovine brains using a modified version of the high-PIPES method (Castoldi and Popov, 2003), wherein the first polymerization cycle was performed in 100 mM PIPES instead of 1 M PIPES. Tetramethylerhodamine (TAMRA) and Alexa Fluor 546 succinimidyl esters were used to label ε-amines on tubulin (Hyman et al., 1991). The labeling reaction is performed on polymerized tubulin so as not to modify the polymerization interfaces. GMPCPP stabilized microtubules were prepared as follows: A polymerization mixture was prepared with BRB80 (80 mM PIPES-KOH, pH 6.9, 1 mM EGTA, 1 mM MgCl_2_)+ 2 mM tubulin, 1 mM GMPCPP (Jena Biosciences), 1 mM MgCl_2_ and a 1:4 molar ratio of TAMRA-labelled/unlabelled tubulin. The mixture was incubated on ice for 5 min, followed by incubation at 37 °C for 2 hr. The polymerized GMPCPP microtubules were centrifuged at maximum speed in a Beckman Airfuge and resuspended in BRB80. GMPCPP seeds were prepared by polymerizing a 1:4 molar ratio of TAMRA-labelled/unlabelled tubulin in the presence of guanosine-5′-[(α, β)-methyleno]triphosphate (GMPCPP, Jena Biosciences) in two cycles, as described previously (Gell et al., 2010). GMPCPP seeds prepared in this way were stable for several months at −80 °C.

#### Total internal reflection fluorescence microscopy and preparation of microscope chambers

The microscope set-up uses a Zeiss Axiovert Z1 microscope chassis, a ×100 1.45 NA Plan-apochromat objective lens, and the Zeiss TIRF III slider. A *λ* = 491 nm diode-pumped solid-state laser (Cobolt) was coupled to a fiber optic cable in free space and introduced into the Zeiss slider. Epifluorescence was achieved using a PhotofluorII excitation source (89 North) with wavelength-specific filter cubes (Chroma). Images were recorded using an Andor iXon + DV-897 EMCCD cameras. Microscope chambers were constructed using custom-machined mounts (Gell et al., 2010). In brief, cover glass was cleaned and silanized as described previously (Helenius et al., 2006). Cover glasses (22 × 22 mm and 18 × 18 mm) were separated by two layers of double-sided tape creating a channel for the exchange of solution. Image acquisition was controlled using MetaMorph (Molecular Devices).

Seeds or GMPCPP stabilized microtubules depending on the experiment were adhered to silanized glass slides as described previously (Bechstedt and Brouhard, 2012). On the day of each experiment for the dynamic assay, aliquots of unlabelled and Alexa Fluor 546-labelled tubulin were thawed, mixed to a 1:3 molar labelling ratio, aliquoted again, and stored in liquid nitrogen.

Microtubule growth from GMPCPP seeds was achieved by incubating flow channels with tubulin in standard polymerization buffer: BRB80, 1 mM GTP, 0.1 mg ml^−1^ BSA, 1% 2-mercaptoethanol, 250 nM glucose oxidase, 64 nM catalase, 40 mM D-glucose. Assays were performed with an objective heater set to 35 °C. Time-lapse image sequences were acquired at 5 s intervals. MCAK depolymerization of GMPCPP stabilized seeds was achieved by incubating flow channels with MCAK and standard polymerization buffer plus ATP: BRB80, 1 mM GTP, 0.1 mg ml^−1^ BSA, 1% 2-mercaptoethanol, 250 nM glucose oxidase, 64 nM catalase, 40 mM D-glucose and 1mM ATP. Assays were performed with an objective heater set to 35 °C. Time-lapse image sequences were acquired at 5 s intervals.

#### Microtubule growth and shrinkage rates

Microtubule growth rates were analyzed by manually fitting lines to kymographs of growing microtubules using the Kymograph and Linescan features in MetaMorph (Molecular Devices).

### Fluorescence in situ hybridization

HCT116 and HCT116 p53^-/-^ cells grown on 22 mm coverslips were washed once with 1× PBS and fixed with (3:1) methanol-acetic acid solution at room temperature for 30 minutes. Cells on coverslips were dried at room temperature for 10 minutes and immersed in 2X SSC (0.3 M NaCl, 30 mM sodium citrate) for 5 minutes at room temperature. Cell were dehydrated in ethanol series (70%, 85%, 100%) each for 5 minutes at room temperature and then air-dried for 10 minutes. Cells were processed for FISH by using specific α-satellite probes against chromosome 7 and 11 (Cytocell) according to the manufacturers protocol. Coverslips were mounted using ProLong gold with DAPI overnight at room temperature and sealed. Chromosome signals from approximately 1000 cells were scored using the criteria outlined in (Van Stedum and King, 2002).

### Statistical analysis

Statistical significance was determined by performing two-tailed *Student’s* t-test with unequal-variance unless otherwise stated. Statistical analysis of all karyotype studies, quantification of lagging chromosomes in *GTSE1*-over expressing clonal cell lines, and phenotype quantification in *GTSE1* knockout clonal cell lines were performed using chi-squared tests.

## Supplemental Material

Figure S1 contains western blots showing RNAi depletion efficiency for all siRNAs and cell lines analyzed. Figure S2 contains verification and characterization of Cas9 nuclease-mediated GTSE1-GFP and GTSE1 knockout clones. Figure S3 includes analysis of kinetochore-MT stability, TACC3 localization, and Kif2A dependence after perturbation of GTSE1 expression. Figure S4 shows purified GTSE1 and MCAK proteins used and in vitro analysis of GTSE1 inhibition of MCAK. Figure S5 contains quantification of different categories of anaphase chromosome segregation defects in U2OS and HeLa cells after GTSE1 and/or MCAK RNAi depletion. Videos 1 and 2 show chromosome dynamics and mitotic progression in control (Video 1) and GTSE1-depleted (Video 2) U2OS cells. Videos 3 and 4 show GTSE1 localization to kinetochore-microtubules in Z-sections from a cold-treated fixed cell labeled by CREST antibody and GTSE1 (Video 3) and tubulin (Video 4). Videos 5 and 6 show MCAK activity on GMPCPP stabilized microtubules with MCAK alone (Video 5) or MCAK and GTSE1 (Video 6).

## Acknowledgements

We thank D. Compton (Dartmouth Medical School) for providing the U2OS PA-GFP-tubulin and HCT116 p53^-/-^ cell lines. We thank A. Musacchio and G. Vader for comments on the manuscript. We thank S. Maffini for help with performing and analyzing photoactivation experiments, A. Faesen for advice and help with microtubule sedimentation assays, M. Mattiuzzo for help with FISH, and J. Beermann and K. Klare for help with protein purification. This work was supported by the Max Planck Institute of Molecular Physiology, a Worldwide Cancer Research project grant to A.B., and an IMPRS-CMB PhD fellowship to S.B. The authors declare no competing financial interests.

## Author Contributions

Shweta Bendre, Conrad Hall, Arnaud Rondelet, Gary Brouhard, and Alexander Bird conceived and designed experiments.

Shweta Bendre and Arnaud Rondelet performed all cellular assays, image quantification, and analysis.

Shweta Bendre performed all protein-interaction studies and GTSE1 purification.

Conrad Hall performed TIRF imaging experiments and analysis.

Arnaud Rondelet designed, created, verified, and characterized Cas9/CRISPR knockout cell lines.

Nadine Wöstehoff created Cas9/CRISPR knockout cell lines

Yu-Chih Lin created GST-fusion constructs.

Shweta Bendre and Alexander Bird wrote the manuscript.

## Abbreviation List

**BAC** Bacterial artificial chromosome

**CIN** Chromosomal instability

**FISH** Fluorescence in situ hybridization

**MT** Microtubule

**TIRF** Total internal reflection fluorescence

## Figure Legends

**Figure S1.** Western blots showing RNAi depletion efficiency for all siRNAs and celllines analyzed. (A) Western blot of cell lysates following control (with dilutions indicated) or GTSE1 RNAi in U2OS cells, probed with anti-GTSE1 and anti-alpha-tubulin. GTSE1 levels are depleted to less than 10% (B) Western blot of cell lysates following control or GTSE1 RNAi in U2OS cells, two stable U2OS cell clones expressing RNAi-resistant GTSE1 (GTSE1 ^WT(204)^ and GTSE1 ^WT(212)^), and two stable U2OS cell clones expressing RNAi-resistant GTSE1 mutated to abolish interaction with EB1 (GTSE1 ^Sk(202)^ and GTSE1 ^Sk(208)^). Blots were probed with anti-GTSE1 and anti-alpha-tubulin. (C) Western blot of cell lysates following control (with dilutions indicated) or GTSE1 RNAi in U2OS H2A.Z-mCherry cells, probed with anti-GTSE1 and anti-alpha-tubulin. GTSE1 levels are depleted to less than 25%. (D) Western blot of cell lysates following control (with dilutions indicated) or GTSE1 RNAi inU2OS PA-GFP-tubulin cells, probed with anti-GTSE1 and anti-alpha-tubulin. GTSE1 levels are depleted to less than 20%. (E) Western blot of cell lysates following control (with dilutions indicated) or MCAK RNAi in U2OS cells, probed with anti-MCAK and anti-alpha-tubulin. MCAK levels are depleted to less than 20%. (F)Western blot of cell lysates following control (with dilutions indicated) or GTSE1 and MCAK RNAi in U2OS cells, probed with anti-GTSE1, anti-MCAK, and anti-alpha-tubulin. GTSE1 and MCAK levels are both depleted to less than 10%. (G)Western blot of cell lysates following control, GTSE1, Kif2A RNAi and co-depletion of Kif2A and GTSE1 in U2OS cells, probed with anti-GTSE1, anti-Kif2A, and anti-alpha-tubulin antibodies. (H) Western blot of cell lysates following control or GTSE1 RNAi in U2OS cells and Hela cells expressing BAC based hKif2C-LAP, probed with anti-GTSE1, anti-MCAK, and anti-alpha-tubulin antibodies.

**Figure S2.** Verification and characterization of Cas9 nuclease-mediated GTSE1-GFP and GTSE1 knockout clones. (A) Western blots showing U2OS cells with endogenous GTSE1 protein tagged with GFP via Cas9-mediated homologous recombination. (B)Western blot of cell lysates from U2OS cells and two Cas9-mediated GTSE1 knockout U2OS clones run by SDS-PAGE and probed with antibodies against GTSE1, MCAK and alpha-tubulin. (C) Scheme representing the GTSE1 locus and the two chosen sites targeted by the Cas9 nuclease for knockouts. The sequence targeted by each Cas9/guide RNA pair is indicated. (D) Regions surrounding the CRISPR-Cas9 cutting sites were amplified by PCR from genomic DNA, sequenced, and heterozygous insertions and deletions were resolved using TIDE software. Histograms show the percentage of sequences showing no modification (n=0), an insertion (n>0) or a deletion of n nucleotides (n<0) at the CRISPR-Cas9 cutting site in exon 2 and exon 9 of the GTSE1 locus. Clones were generated by transfecting U2OS cells with a Cas9 nuclease targeting both exon 2 and 9 (Clone 1) or exon 9 alone (Clone 2), respectively. (E) Quantification of spindle length in U2OS cells following control and GTSE1 RNAi (n ≥ 145 cells per condition over 4 independent experiment) and U2OS GTSE1 ^KO(1)^ and GTSE1 ^KO(2)^ cells lines (n ≥ 50 cells each). (F-H) Western blot of cell lysates from U2OS cells and two Cas9-mediated GTSE1 knockout U2OS clones run by SDS-PAGE and probed with antibodies against pS558-TACC3, TACC3, ch-Tog, Aurora B and alpha-tubulin. (I) Immunofluorescence images of mitotic GTSE1 knockout clones after control and MCAK RNAi, stained for DNA (DAPI) and microtubules (tubulin). Scale bar represents 5 microns.

**Figure S3.** Analysis of kinetochore-MT stability, TACC3 localization, and Kif2A dependence after perturbation of GTSE1 expression. A) Graph showing average normalized fluorescence intensity at each time point following photoactivation of PA-GFP-tubulin in U2OS metaphase spindles treated with control or GTSE1 RNAi. n = 11 cells per condition over three independent experiments. Solid lines represent the double exponential fit for control and GTSE1 RNAi. Error bars represent standard error of the mean. (B) Immunofluorescence images of mitotic U2OS cells following control and GTSE1 RNAi and stable GTSE1 knockout clones, stained for microtubules (tubulin) and TACC3. Bar chart represents quantification of TACC3 on mitotic spindle in U2OS cells following control or GTSE1 RNAi (n ≥ 50 cells per condition over 3 independent experiments) and in stable GTSE1 knockout clones (n ≥ 50 cells each over 2 independent experiments). Error bars represent standard error of the mean.Differences were not statistically significant. (C) Immunofluorescence images of mitotic U2OS cells following control, GTSE1, Kif2A, or combined GTSE1 and Kif2A RNAi, stained for DNA (DAPI), Kif2A, and microtubules (tubulin). (D)Immunofluorescence images of mitotic HCT116 control and GTSE1-GFP overexpressing clones stained for microtubules (tubulin) and TACC3. Graph represents average tubulin fluorescence of the inner spindle from immunofluorescence analysis of HCT116 control and GTSE1-GFP overexpressing clones. Second graph shows average TACC3 intensity of the inner spindle quantified from immunofluorescence analysis of HCT116 control and GTSE1-GFP overexpressing clones. Third graph represents average TACC3 intensity on the inner spindle normalized to tubulin intensity for HCT116 control and GTSE1-GFP overexpressing clones. n ≥ 52 cells per condition for 1 experiment. P-values were obtained using an Anova and Tukey’s test. Error bars represent standard deviation. Scale bars represent 5 micrometers.

**Figure S4.** In vitro analysis of GTSE1 inhibition of MCAK. (A) Coomassie-stained SDS-PAGE gels of inputs from in vitro GST pull-down assays. (B) Gel showing full length MCAK and GTSE1 purified from baculovirus-infected insect cells. Molecular weights are indicated to the left. (C) Size exclusion chromatography (SEC) profile of FL-GTSE1 (78.5 kDa) shows that the protein elutes earlier than expected, consistent with an elongated unfolded structure. (D) SEC profile of FL-MCAK shows that MCAK (81 kDa) runs as a dimer. (E) In vitro sedimentation assay to monitor inhibition of MCAK depolymerase activity by GTSE1. Taxol stabilized microtubules were added to 200 nM MCAK and varying concentrations of GTSE1 in presence of ATP. The reaction mixture was incubated for 1 h at room temperature followed by centrifugation at high speed. Supernatants and pellets were collected and equal amounts were run on SDS-PAGE gel and stained with Coomassie blue. The histogram shows the percentage of tubulin for each condition in the pellet fraction, quantified from the density of the Coomassie gel from a representative experiment.

**Figure S5.** Quantification of anaphase chromosome segregation defects in U2OS and HeLa cells after GTSE1 and/or MCAK RNAi depletion. (A) Quantification of anaphase lagging chromosomes. (B) Quantification of chromosome bridges. (C)Quantification of acentric fragments. Error bars represent standard error of the mean. n > 220 U2OS cells with defective anaphases over 3 independent experiments. n > 390 HeLa cells with defective anaphases over 3 independent experiments. p≤0.05 *, p≤0.01 **, p≤0.001 ***

### Supplemental Video Legends

**Video 1. Chromosome dynamics and mitotic progression in control U2OS cells.** Mitotic cells efficiently align chromosomes and enter anaphase. U2OShis tone H2A.Z-mCherry cells after control RNAi were imaged at 1 minute intervals,and displayed at 8 frames per second.

**Video 2. Chromosome dynamics and mitotic progression in GTSE1-depleted U2OS cells.** Mitotic cells have difficulty aligning chromosomes and delayed mitotic timing, yet are able to eventually align chromosomes and enter anaphase. U2OShis tone H2A.Z-mCherry cells after GTSE1 RNAi were imaged at 1 minute intervals,and displayed at 8 frames per second.

**Video 3. GTSE1 localization to kinetochore microtubules**. Representative Z-stack of a U2OS cell showing GTSE1 localization to kinetochore microtubules. The U2OS cells were treated with cold for 17 minutes and then stained for GTSE1 and CREST (kinetochore). Movie is displayed at 8 frames per second.

**Video 4. Kinetochore microtubules in U2OS cells.** Representative Z-stack of a U2OS cells (same as in Movie 3) showing microtubules in grey (tubulin) and kinetochores (CREST) in red. Movie is displayed at 8 frames per second.

**Video 5. GMPCPP stabilized microtubules in the presence of 50 nM MCAK.** Epifluorescence images of GMPCPP stabilized microtubules were captured at 5 s intervals for 5 minutes. The GMPCPP stabilized microtubules were observed to depolymerize rapidly from both ends in the presence of 50nM MCAK. Video playback is 45× real-time (see time stamp).

**Video 6. GMPCPP stabilized microtubules in the presence of 50 nM MCAK and 250 nM GTSE1.** Epifluorescence images of GMPCPP stabilized microtubules were captured at 5 s intervals for 5 minutes. The GMPCPP stabilized microtubules were observed to maintain a constant length in the presence of 50 nM MCAK and 250 nM GTSE1. Video playback is 45× real-time (see time stamp).

